# Diurnal regulation of Acyl-CoA synthetase 3 (ACSF3) underlies daily mitochondrial lysine-malonylation and hepatic metabolism

**DOI:** 10.1101/2024.09.03.607283

**Authors:** Enora Le Questel, Charlène Besnard, Florian Atger, Yolène Foucher, Alwéna Tollec, Victoria Pakulska, Arsênio Rodrigues Oliveira, Chloé Clotteau, Mathilde Gourdel, Ivan Nemazanyy, Mikael Croyal, Yohann Coute, David Jacobi, Bertrand Cariou, Daniel Mauvoisin

**Author notes:** F.A. is currently employed by Novo Nordisk Foundation Center for Basic Metabolic Research, Faculty of Health and Medical Sciences, University of Copenhagen, Denmark. Y.F. is currently employed by Namsa, Chasse-Sur-Rhone, France.

## Abstract

Circadian rhythms are fundamental to maintaining health and are implicated in various diseases. In the liver, daily rhythms are coordinated via the interplay between feeding rhythms and the molecular circadian clock, ensuring metabolic homeostasis. Disruption of feeding rhythms can lead to circadian misalignment, contributing to metabolic disorders, yet the underlying molecular mechanisms remain unclear. Recent evidence suggests that post-translational modifications play a key role in regulating circadian functional output. In this framework, mitochondria serve as a convergence point, integrating rhythms in metabolism, feeding rhythms and the circadian clock. In the present study, we used a multi-omics approach to investigate the role of the Acyl-CoA synthetase 3 (ACSF3) in driving lysine-malonylation and in regulating daily hepatic metabolism. We found that ACSF3 expression and its mediated impact on lysine-malonylation are rhythmic and largely governed by feeding rhythms. While hepatic ACSF3 knockdown did not alter diet-induced metabolic abnormalities, our results demonstrate that ACSF3 plays a role in the diurnal regulation of liver glycogen storage, *de novo* lipogenesis, and triglyceride synthesis.

## INTRODUCTION

A modern lifestyle, which often includes chrono-disruptive habits like irregular feeding patterns, disrupts our daily rhythms and contributes to metabolic pathologies such as type 2 diabetes^1^. Daily rhythms are coordinated by the interplay between both environmental factors such as feeding rhythms and the molecular circadian clock, allowing metabolic homeostasis^2^. In fact, circadian gene expression regulation occurs at several regulatory steps ranging from transcription^3,4^ to protein accumulation^5,6^ and post-translational modifications (PTMs)^7–9^, the latter playing a major role in driving circadian functional outputs^10^. To date, the link between biological rhythms and metabolic disorders remains poorly understood and characterizing the molecular regulatory nodes between daily rhythms and metabolism is utmost.

In this framework, mitochondria appear as a key hub allowing metabolic rhythmicity integration in response to circadian and feeding rhythms. Indeed, mitochondrial rhythmicity is evidenced at the transcriptional^11^, post-transcriptional^12^ and post-translational levels^9,13^ contributing to the diurnal regulation of key processes such as oxidative phosphorylation^14,15^, calcium homeostasis^16,17^ or mitochondrial dynamics^13,18^. In fact, mitochondria acidic environment favours spontaneous PTMs occurrence such as lysine-acetylation from acetyl-CoA^19^. In liver mitochondria, lysine-acetylation rhythmicity is mainly controlled by Sirtuin-3 (SIRT3) dependant deacetylation processes driven by NAD^+^ rhythmicity^9,20^. Among the Sirtuins expressed in mitochondria, SIRT5 has a weak deacetylase and higher demalonylase activity^21,22^. Lysine-malonylation relies on malonyl-CoA availability, a metabolite playing a crucial role in *de novo* lipogenesis and in β-oxydation regulation^23^. Proteomics work using *Sirt5* Knockout (KO) mice has uncovered the link between malonylation and glycolysis regulation^24^. SIRT5 hepatic overexpression improves the metabolic phenotype of genetic obese mouse model^25^, suggesting a role of decreased hepatic lysine-malonylation in metabolic disorders.

Using malonate as the main substrate, Acyl-CoA synthetase 3 (ACSF3) produces mitochondrial malonyl-CoA and fuels the TCA cycle via the malonyl-CoA decarboxylase (MLYCD)^26^ or type II fatty acid synthesis^27^. While the ACSF3 substrate impairs oxidative phosphorylation^28^, its expression controls mitochondrial lysine-malonylation^29^.

Although disruption of ACSF3 expression *in vitro* has been linked to metabolic disturbances^28^, knowledge on the role of lysine-malonylation and ACSF3 in daily liver metabolism is scarce. While hepatic malonyl-CoA level is rhythmic and controlled by feeding^30^, hepatic malonylation levels are globally disturbed in circadian clock deficient *Bmal1* or *Cry1/Cry2* KO mice^9^. Moreover, ACSF3 has been shown to impact the circadian clock period via post-transcriptional mechanisms^31^. These observations suggest that ACSF3 modulated malonylation could be a regulatory node of daily hepatic metabolism.

In the present work, we first investigated the diurnal regulation and the role of hepatic ACSF3 and its mediated lysine-malonylation in maintaining hepatic metabolic homeostasis. First, we examined the diurnal regulation and the role of the circadian clock and feeding schedule on the expression of ACSF3 and other targets that could impact lysine-malonylation. We also studied the downstream effect on global and mitochondrial lysine-malonylation. Second, we analyzed the consequences of inhibiting hepatic ACSF3 expression on mouse metabolic outcome. This experiment led us to characterize the impact of hepatic *Acsf3* Knock-Down (KD) on the malonyl-proteome (malonylome), the diurnal metabolome as well as diurnal lipidome signature in mouse liver. In addition to the impairment of oxidative phosphorylation, our observations support a role of hepatic ACSF3 and its mediated effect on lysine-malonylation in finely tuning diurnal glycogen store mobilization, *de novo* lipogenesis and triglycerides (TG) synthesis.

## RESULTS

### Regulation of total versus mitochondrial proteome uncovers ACSF3 and lysine-malonylation diurnal rhythmicity

To get further insights into the diurnal regulation of the mitochondrial proteome, we first analyzed in parallel the rhythmicity of proteomic data in mouse liver at the whole tissue^32^ vs. the mitochondrial level^12^ using CircaCompare^33^ (see Methods). Like previous work, ∼5% and 25% of the 1650 proteins quantified in both studies were rhythmic in total extract (TE) and mitochondrial extract (ME) respectively (Fig. S1a; Table S1). For those proteins exclusively rhythmic in ME, 175 exhibited differences in mean level in circadian clock deficient mouse liver (*Bmal1* KO or *Cry1/Cry2* KO)^32^ (Fig. S1b; Table S1). Gene enrichment analysis using MitoCarta 3.0 pathways^34^ revealed that, rhythmic proteins in ME that are unaltered in circadian clock deficient animals are enriched in pathways regulated by calcium^35^ and type II fatty acid synthesis (Fig. S1c; Table S1). Initiated by ACSF3, type II fatty acid synthesis uses discrete monofunctional enzymes like malonyl-CoA-ACP acyltransferase (MCAT) and mitochondrial trans-2-enoyl-CoA reductase (MECR) to eventually produce lipoic acid^27^.

Hence, using western blot (WB), we monitored ACSF3, MCAT and MECR, in liver TE and ME collected every 3h over 24h from mice subjected to nighttime (mouse activity period) restricted feeding during four days (Fig. 1a) as in Mauvoisin *et al.* 2014^5^. This feeding condition is known to increase the amplitude of liver weight rhythmicity as already described^36^ and confirmed here (Fig. S1d). The ACSF3 diurnal rhythm profile was highly pronounced in ME with a peak phase at ZT21 (Fig. 1b). MCAT and MECR were also rhythmic in ME peaking at the beginning of the day (Fig. 1b). Malonylation level was rhythmic in ME peaking a bit later than ACSF3 (Fig. 1c). SIRT5 and MLYCD expression were rhythmic in ME with a peak phase at ZT3 and ZT18, respectively (Fig. 1b). Using targeted metabolomics, we quantified the diurnal rhythm of ACSF3 substrates, malonate and methyl-malonate, and observed a peak at ZT0 in the liver (Fig. S1e). As a proxy of type II fatty acid synthesis, we also monitored lipoylation level. Overall, the lipoic acid signal was not significantly rhythmic in TE or ME (Fig. 1d).

**Figure 1.**
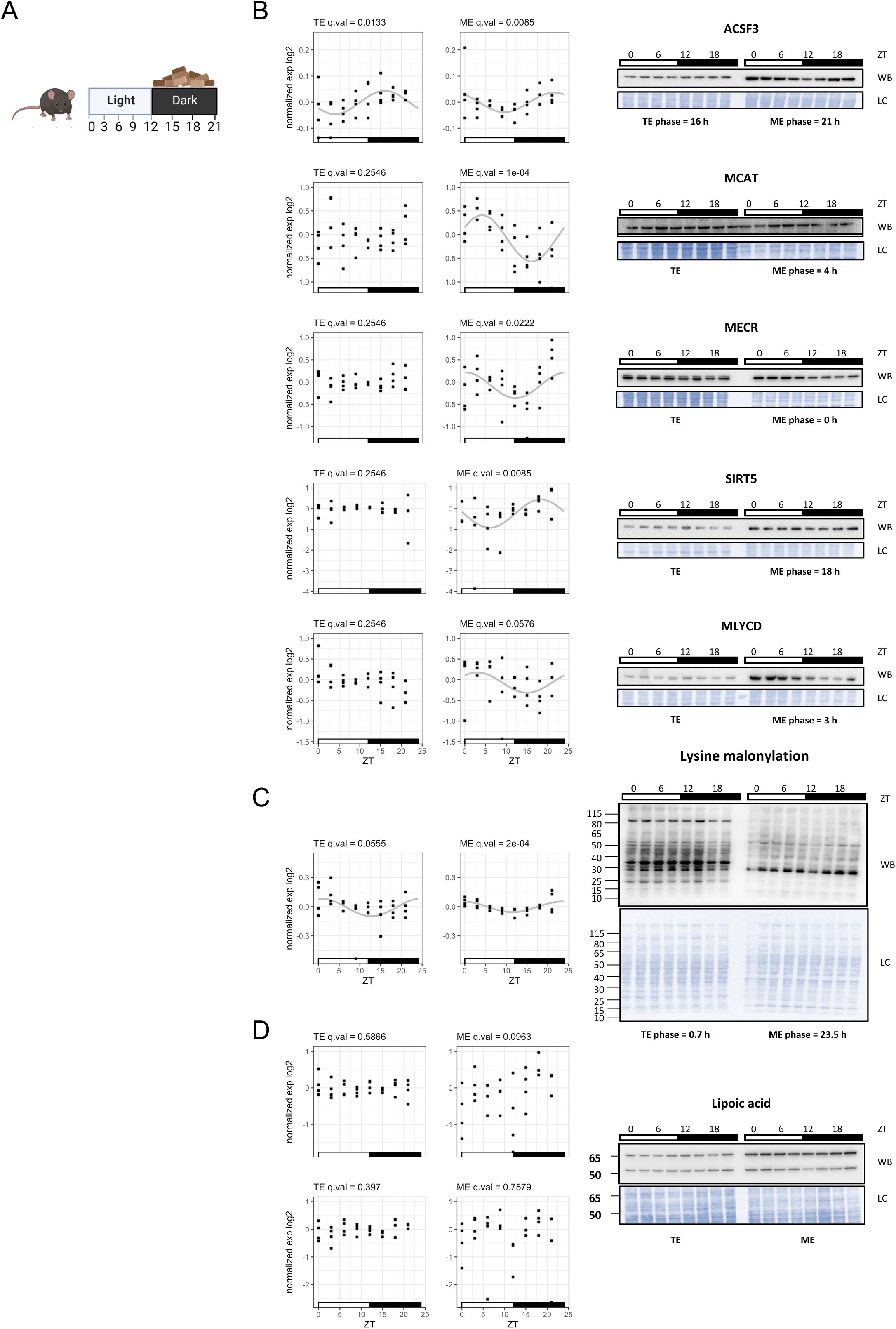
**a.** Experimental design. Four days of night-restricted feeding was performed in mice fed a CD. Livers were collected at 3-hour intervals (n=4 days). **b-d.** Western blot (WB) analysis of **(b)** ACSF3, MCAT, MECR, SIRT5, MLYCD, **(c)** lysine malonylation and **(d)** lipoic acid level in total extracts (TE) and mitochondrial extracts (ME) over time. The graphs display densitometry analysis of all replicates. LC: loading control. Data are normalized by the temporal mean in each extract. Grey lines correspond to the CircaCompare fit displayed if q.val < 0.05. In **(d)**, the upper and lower graphs are respectively the quantification of the 65 and 50 kDa bands.

To get more information about the role of the molecular circadian clock in regulating other type II fatty acid synthesis and malonylation regulators diurnal expression, we analyzed the rhythmicity of hepatic RNAseq data from *Bmal1* KO and *Cry1/2* KO mouse housed in NRF during four days^37^ (Table S2). With the exception of *Sirt5* in *Cry1/2* WT and *Acsf3* and *Mlycd* in *Bmal1* WT, no significant rhythmicity of type II fatty acid synthesis or mitochondrial malonylation regulator transcripts was observed in WT or circadian clock deficient mouse liver (Fig. S1f;Table S2). Proteomic data from Mauvoisin *et al.* 2014^5^ (Table S1), showed that type II fatty acid synthesis and malonylation regulator protein levels were not significantly disturbed in circadian clock deficient mouse liver (Fig. S1g; Table S1).

Altogether, these data showed a pronounced rhythmicity of ACSF3 expression, followed by malonate and malonylation in ME. These data also suggest that this daily rhythm is influenced by feeding rather than by the molecular circadian clock.

### Feeding schedule regulates hepatic diurnal ACSF3 expression and impact mitochondrial lysine-malonylation

To study the impact of feeding schedule, we analyzed the rhythmicity of RNAseq from (GSE118967) and (GSE159135) who studied the impact of night feeding (nRF)^38^ or day feeding (dRF)^39^ vs. *Ad libitum* (AL) in mouse liver during five and four weeks, respectively. These data showed that nRF tended to imprint diurnal rhythmic expression of *Acsf3*, *Mcat*, and *Mlycd* transcripts, while dRF reversed the phase of the rhythmic expression of *Acsf3* and *Mlycd* (Fig. S2a; Table S2).

To study this regulation at the protein level, we subjected mice to the same feeding paradigms during a week (AL, nRF, dRF) and monitored by WB the expression of type II fatty acid synthesis proteins as well as malonylation level and its regulators at ZT0 vs. ZT12 (Fig. 2a). These feeding paradigms are known to impact liver weight rhythmicity. We observed that nRF reinforce ZT0 vs. ZT12 difference in liver weight compared to AL, while dRF reversed it, but with a lower amplitude (Fig. S2b). ACSF3 was also impacted by feeding schedule in ME. Indeed, although the difference was not significant in AL, ACSF3 level was higher at ZT0 vs. ZT12 in nRF and this temporal difference tended to be reversed in dRF (Fig. 2b; Fig. S2c). MCAT and MECR levels were also impacted by feeding schedule in TE but not in ME, while SIRT5 and MLYCD expressions were not significantly modified (Fig. 2b; Fig. S2c).

**Figure 2.**
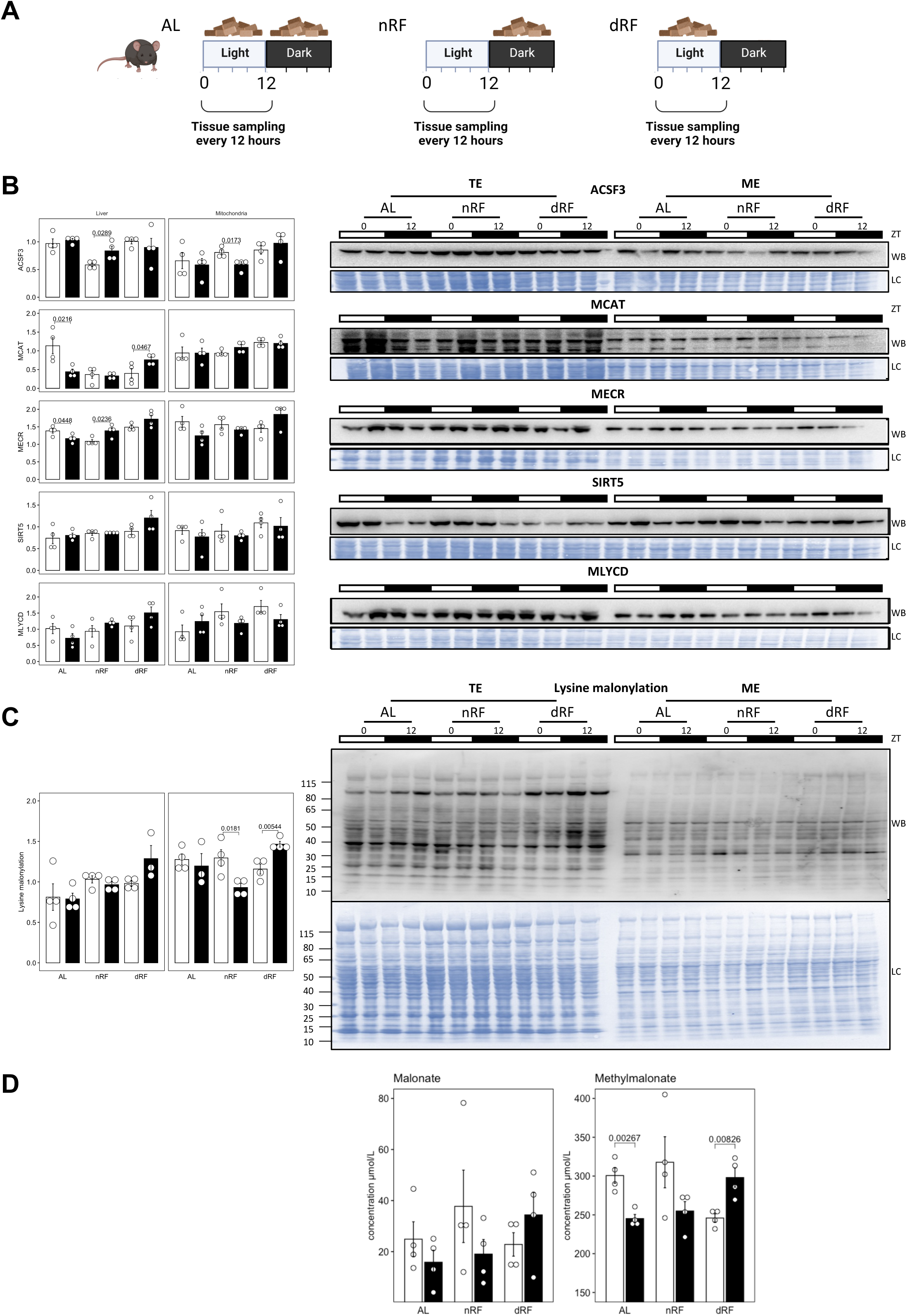
**a.** Experimental design. *Ad libitum* (AL) vs. night-restricted feeding (nRF) vs. day-restricted feeding (dRF) was performed in mice fed a CD for a week. Livers were collected at ZT0 and 12 (n=4 days). **b-c.** Western blot (WB) analysis of **(b)** ACSF3, MCAT, MECR, SIRT5, MLYCD, **(c)** lysine malonylation and in total extracts (TE) and mitochondrial extracts (ME) (n=2 replicates) over time. The graphs display densitometry analysis of all 4 replicates displayed in FigS2c for ACSF3, MCAT, MECR, SIRT5 and MLYCD and FigS2d for lysine malonylation. LC: loading control. For each extract, ZT0 vs ZT12 were compared using Tukey HSD test. **d.** Liver malonate (left panel) and methylmalonate (right panel) levels over time (Data presented as mean ± SEM). ZT0 vs ZT12 were compared using Tukey HSD test.

Similarly to ACSF3, malonylation level was higher at ZT0 compared to ZT12 upon nRF in ME and this difference was reversed by dRF (Fig. 2c; Fig. S2d). On the other hand, lipoic acid signal was not modified by feeding schedule in ME (Fig. S2e). Although not significantly modified, ACSF3 substrate levels followed similar pattern as ACSF3 expression did (Fig. 2d). Together, these observations showed that ACSF3 and its mediated malonylation observed in ME are regulated by feeding behavior.

### Diet induced obesity modifies ACSF3 diurnal rhythmic expression

By modifying rhythmic food intake, diet induced obesity (DIO) strongly impacts daily hepatic metabolism^40–42^. To study the impact of DIO on type II fatty acid synthesis and lysine-malonylation regulators, we first analyzed RNAseq data from mice subjected to 12 weeks of High Fat Diet (HFD)^43^ (GSE108688). While HFD led to an increase of *Acsf3*, *Mcat*, *Mecr* and *Sirt5* expression, we observed an increased diurnal rhythmicity for transcripts encoding proteins implicated in malonylation regulation *i.e. Acsf3*, *Sirt5* and *Mlycd* (Fig. S3a; Table S2). To get more information at the protein level, we induced obesity in mice by feeding them a HFD during 12 weeks (Fig .3a), collected liver samples every 3h and analyzed protein levels by WB. HFD significantly increased body weight (Fig. S3b) while impairing liver weight rhythmicity (Fig. S3c). In TE, the malonylation level was decreased upon HFD (Fig.3b). Since, malonyl-CoA availability affects lysine-malonylation, this could be associated with the observed increased in malonyl-CoA consuming enzyme MLYCD expression (Fig.3c) and the decrease in *de novo* lipogenesis enzyme Acetyl-CoA Carboxylase Alpha (ACACA) and Fatty Acid Synthase (FASN) in TE (Fig.S3d)^44^. In ME, we observed that diurnal malonylation rhythmicity is increased by HFD, as well as the global level and the rhythmicity of ACSF3 and SIRT5 (Fig.3c). Although, hepatic level of malonate and methylmalonate were not significantly rhythmic, there was a trend for a decrease level of both metabolites in HFD at the night/day transition suggesting a differential diurnal profile upon HFD (Fig.S3e). Lipoylation rhythmicity was also increased by HFD in TE and ME (Fig.S3f).

**Figure 3.**
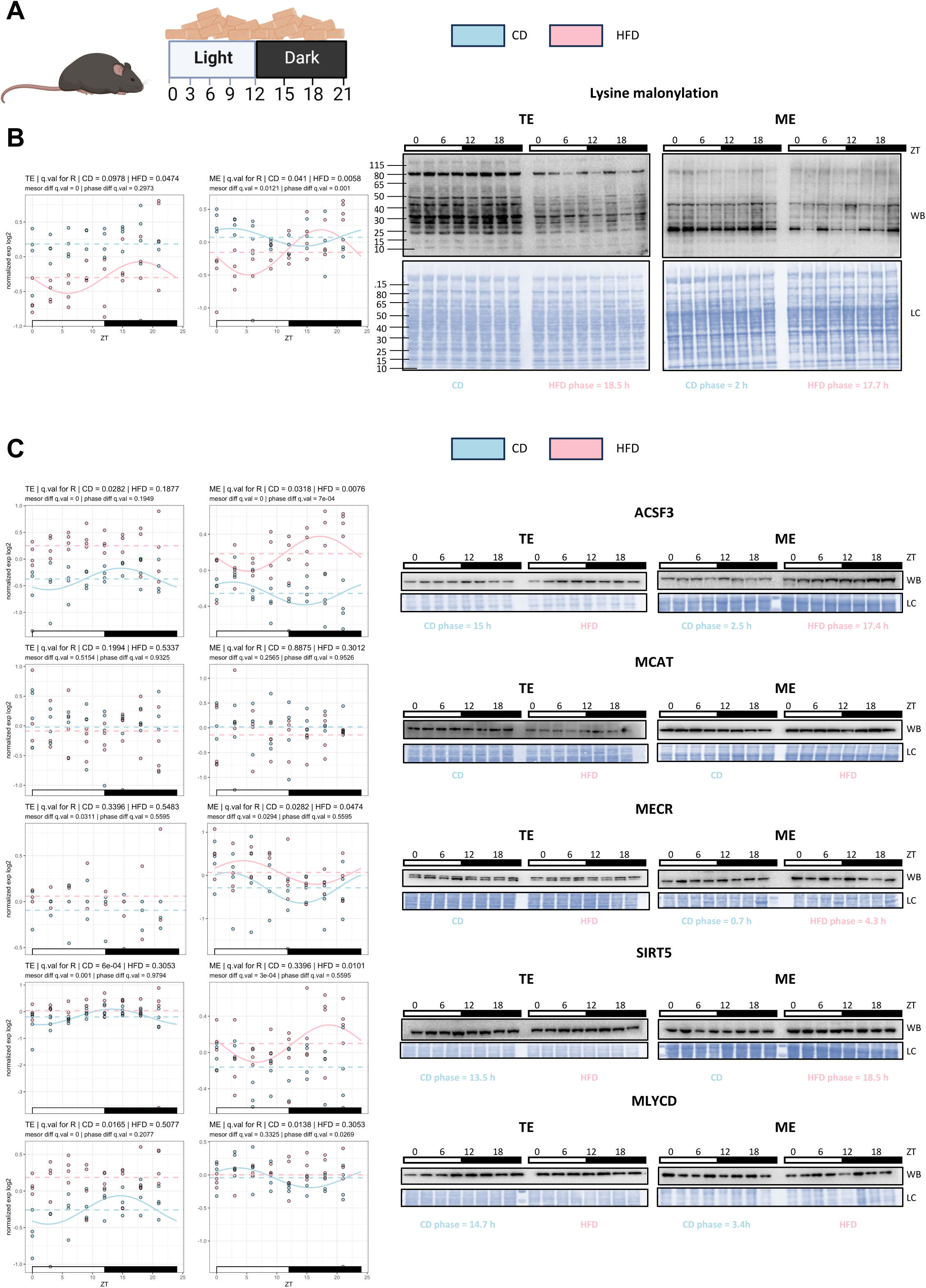
**a.** Experimental design. *Ad libitum* (AL) vs. High Fat Diet (HFD) was performed in mice for 12 weeks. Livers were collected at 3-hour intervals (n=4 days). **b-c.** Western blot (WB) analysis of **(b)** lysine malonylation and **(c)** ACSF3, MCAT, MECR, SIRT5, MLYCD in total extracts (TE) and mitochondrial extracts (ME) over time upon CD (light blue) or HFD (pink). The graphs represent densitometry analysis of all replicates. LC: loading control. Data are normalized to the global mean in each extract. Lines correspond to the rhythmicity fit displayed if q.val < 0.05. Horizontal dashed lines show the mesor levels for each diet. Statistics for differential rhythmicity are also reported in the graph titles.

These results showed that diurnal mitochondrial malonylation is also disturbed upon DIO and this perturbation is associated with the modification of the diurnal expression of lysine-malonylation regulators (*i.e.* ACSF3 and SIRT5).

### Hepatic ACSF3 knockdown impacts glucose metabolism upon control diet

SIRT5 has been shown to regulate hepatic metabolism via controlling lysine-malonylation level of glycolysis proteins such as the lysine 184 of GAPDH during fasting^24,26^. Thus, we next aimed at uncovering the role of hepatic ACSF3 on mice metabolism. To do so, we monitored the metabolic outcomes in mice with a hepatic ACSF3 KD fed a CD vs. a HFD during five weeks (Fig. 4a), which is sufficient to highly increase body weight compared to CD (Fig. S3b). Hepatic ACSF3 KD was performed with shRNA using a AAV delivery strategy (Fig. 4b). Upon CD, ACSF3 KD led to an increase in body weight (Fig. 4c) with no perturbation of the rhythmic (Fig. 4d) or total (Fig. 4e) food intake. Monitoring glycemia during the experiment revealed ACSF3 KD increased glycemia at ZT0 after 4 weeks (Fig. 4f) while decreasing 6-hours-fasting glycemia at ZT6 (Fig. 4g). Meanwhile, glucose tolerance (Fig. 4h) and insulin sensitivity (Fig. 4i) were respectively improved and impaired upon ACSF3 KD. Surprisingly, the aforementioned effects were not present upon HFD (Fig. S4a to Fig. S4g). While, these results show that hepatic ACSF3 KD did not lead to overt metabolic abnormalities under HFD, the observed impact observed under CD is intriguing. Interestingly, knocking out the demalonylase *Sirt5* did not modify metabolic abnormalities under HFD but had slightly opposite effect on insulin sensitivity upon CD^45^. The molecular mechanisms behind these opposite insulin tolerance impacts in both models might come from a differential impact on lysine-malonylation.

**Figure 4.**
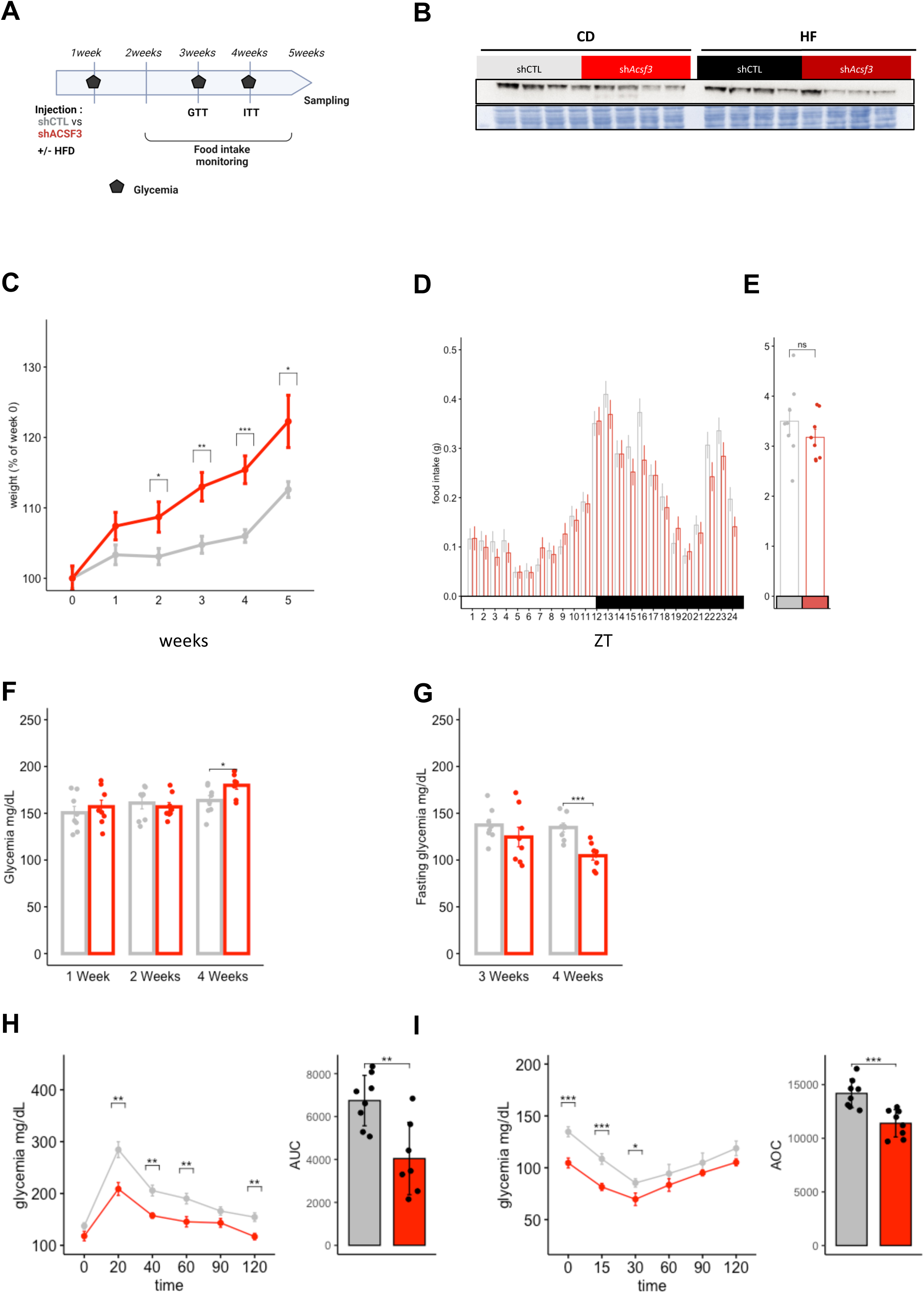
**a.** Experimental design. Hepatic AAV-mediated knockdown (KD) of *Acsf3* was performed in mice subjected to Chow diet (CD) vs High Fat Diet (HFD) for 5 weeks. Glucose Tolerance Test (GTT) and Insulin Tolerance Test (ITT) were performed at 3- and 4-weeks post AAV injection respectively. Food intake was monitored between 2 and 4 weeks. **b.** Validation by Western blot (WB) of AAV-mediated knockdown of ACSF3 in liver at ZT0. **c-i.** Mouse body weight gain (n=16/condition) **(c);** diurnal food intake **(d);** total daily food intake **(e);** glycemia at ZT0 at 1, 2 and 4 weeks **(f);** fasting glycemia after 6 hours of fasting starting at ZT0 at 3 and 4 weeks **(g);** GTT (left) and the corresponding AUC above baseline (right) **(h);** ITT (left) and the corresponding AOC under baseline (right) **(i)** in sh*Ctl* (grey) vs sh*Acsf3 (red)* mice upon CD.

### Hepatic ACSF3 influences malonylation level of proteins implicated in glucose and lipid metabolism

Because upon CD, ACSF3 KD decreased malonylation level in ME (Fig. 5a upper blot), (Fig. 5a lower blot), we next investigated the impact on the global hepatic malonylome and proteome using an *in vivo* Stable Isotope Labeling by Amino acids (SILAC) coupled to mass spectrometry (MS)-based proteomic strategy. After trypsin digestion, protein extracts from five sh*Ctl* and five sh*Acsf3* mouse liver at ZT0 were enriched for malonylated peptides (see Methods and Fig. S5a). Relative abundance of malonylated peptides in the resulting ten TE samples was quantified using a corresponding common SILAC total reference extract (Table S3).

**Figure 5.**
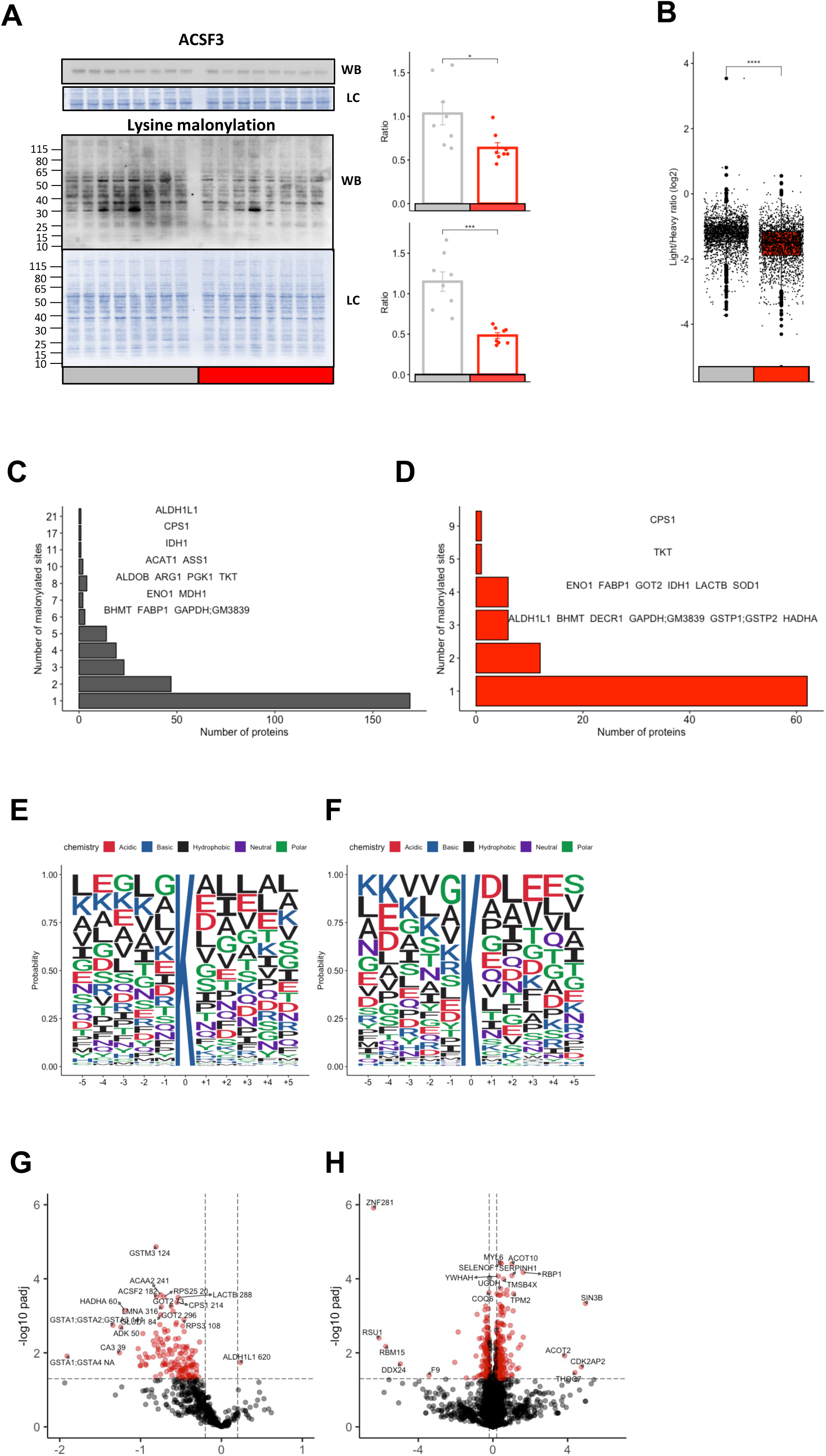
**a.** Western blot (WB) analysis of ACSF3 and malonylation level in liver ME. The left graphs display densitometry analysis of all replicates in sh*Ctl* vs. sh*Acsf3* mice compared using Tukey HSD test. **b.** Global Light/Heavy MS signal of liver malonylation levels normalized by total protein signal in sh*Ctl* vs. sh*Acsf3.* The 5 replicates of sh*Ctl* vs. sh*Acsf3* were compared using Wilcox test. **c-d.** Distribution of the number of malonylation sites identified per protein in all proteins **(c)** vs. in proteins with identified ACSF3-modulated sites **(d)**. **e-f.** Consensus sequence logo plots centered on the lysine of all identified malonylation sites **(e)** or all identified ACSF3 KD-modulated malonylation sites **(f)** (±5 amino acids). **g-h.** Fold change imprinted by ACSF3 KD on the liver malonyl-proteome **(g)** or total liver proteome **(h)** in sh*Ctl* vs. sh*Acsf3* mice compared using Tukey HSD test.

Overall, we identified the malonylation level of 611 sites (Fig. S5a) distributed in 286 proteins. Among them, 365 sites were repeatedly quantified (*i.e.* yielding relative measurements in at least 3 of 5 replicates) (Table S3). Across these malonylated proteins, 169 (59%) had a single malonylation sites and 117 (41%) had two or more sites quantified. The most heavily malonylated proteins were ALDH1L1 and CPS1 with 21 and 17 malonylated sites, respectively (Fig. 5c; Table S3). Both of these proteins remained in the most influenced by ACSF3 KD with 9 and 3 sites significantly modified. Among the other proteins with more than 2 sites impacted by ACSF3 KD, we found. TKT and IDH1 that, which, like ALDH1L1, are implicated in NADPH metabolism. Three other proteins namely ENO1, GAPDH and GOT2 were implicated in glycolysis and gluconeogenesis while two others (DECR1, HADHA) were involved in fatty-acid oxidation (Fig. 5D).

We then compared the frequency of amino acids surrounding all observed malonylated sites using ggseqlogo^46^. We first analyzed data from Nishida *et al.*^24^ in WT versus *Sirt5* KO mice and drew similar conclusion, *i.e.* malonylated sites are surrounded by an over representation of Leucine and to a lower extent Isoleucine at position closed to modified sites enrichment (Fig. S5b) especially at position 2 and -2 in our dataset (Fig. 5e). However, when comparing with the malonylated sites impacted by ACSF3 KD (Fig.5f), we did not observe a diminution of Leucine and Isoleucine proportion, in opposite to what was observed in *Sirt5* KO (Fig.S5c). Differential analysis of ACSF3 KD impact on malonylated lysines coupled with gene Ontology analysis showed a high enrichment of pathways implicated in glucose and lipid metabolism more specifically glycolysis/gluconeogenesis and fatty acid oxidation (Fig. S5d; Fig. S5e). Altogether, these observations indicate no effect of the protein sequence on ACSF3 modulated malonylations. However, the poor overlap between ACSF3 KD impacted vs. *Sirt5* KO impacted malonylation sites (Fig. S5f; 15 (2%) sites among 309 differentially regulated) suggests that ACSF3 regulated malonylome explains the different metabolic phenotype observed between ACSF3 KD and *Sirt5* KO.

### ACSK3 KD disturbs diurnal liver lipid metabolism and promotes TG synthesis

The liver tightly adjusts physiological processes, notably lipids and glucose metabolic pathways to their relevant time of day. Thus, we next aimed at further characterizing the role of hepatic ACSF3 by looking at the impact of its KD on diurnal liver metabolism regulation. We collected samples from sh*Ctl* and sh*Acsf3* mice every three hours over 24h four weeks after AAV injection (Fig. 6a; Fig. 6b) and performed targeted metabolomics. To get more information, we analyzed the rhythmicity of this dataset using CircaCompare (see Methods ; Table S4). Within the 151 metabolites quantified, we first looked at ACSF3 substrate and product and observed that ACSF3 KD significantly increase malonic acid level (Fig. 6c) and decreased the ratio between malonyl-CoA vs. malonic acid while impairing its rhythmicity (Fig. 6d). Proteomics data revealed that ACSF3 KD impacted the expression and the malonylation level of enzymes implicated in NADPH production (Fig. 5d) and, in accordance, we observed that NADPH levels were significantly increased (Fig. 6e).

**Figure 6.**
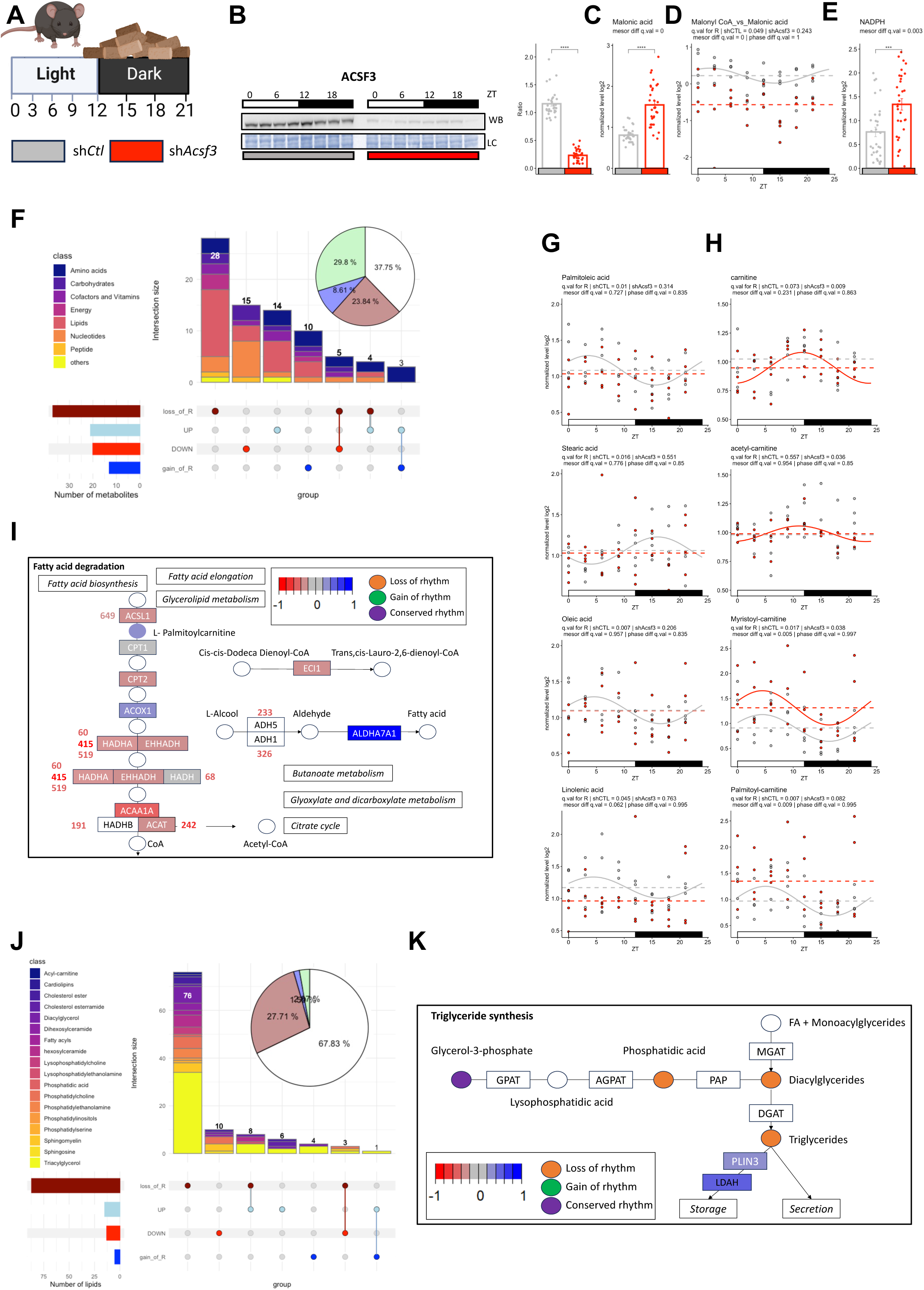
**a.** Experimental design. Hepatic AAV-mediated knockdown (KD) of ACSF3 was performed in mice and subjected to Chow diet (CD) during 4 weeks. Livers were collected at 3-hour intervals (n=4 days). **b.** Western blot (WB) analysis of ACSF3 in liver TE. The left graphs display densitometry analysis of all replicates in sh*Ctl* vs sh*Acsf3* mice compared using Tukey HSD test. **c.** Malonic acid level in TE of sh*Ctl* (grey) vs. sh*Acsf3* (red) mouse liver. The mesors were compared with CircaCompare. **d.** Malonyl-CoA vs. Malonic acid level in TE of sh*Ctl* (grey) vs. sh*Acsf3* (red) mouse liver over time. The graphs display all replicates and dashed lines correspond to mesor level. Rhythmicity statistics and fit (solid line) are also reported. **e.** NADPH level in TE of sh*Ctl* (grey) vs sh*Acsf3* (red) mouse liver. The mesor were compared with CircaCompare. **f.** Matrix layout for all intersections (vertical bars), sorted by size, of differentially regulated metabolites of sh*Ctl* vs. sh*Acsf3* mouse liver between the metabolites that loss or gain rhythmicity or with significant up-regulated (UP) or down-regulated (DOWN) mesor. Color-coded circles in the matrix indicate sets that are part of the intersection and, the top horizontal bar graph displays the metabolites number. Each category of metabolites is color coded. The inset pie plot displays the proportion of metabolites gaining (blue), losing (dark red) or conserving (green) rhythmicity upon ACSF3 KD or non-rhythmic metabolites (white). **g-h.** Fatty acids **(g)** or carnitine and acyl-carnitines **(h)** level in TE of sh*Ctl* (grey) vs. sh*Acsf3* (red) mouse liver over time. The graphs display all replicates and dashed lines correspond to the mesor level. Rhythmicity statistics and fit (solid line) are also reported. **i.** Rhythmic metabolites and differentially regulated proteins at total and malonylated levels in fatty acid degradation pathway (adapted from KEGG pathway database). Each dot represents a class of lipids, with color coding differential levels or changes in rhythmicity. Each rectangle represents a protein and the impact of ACSF3 KD on protein fold change is color coded. The impact on malonylation levels (if present) is indicated by the presence of the position number of the malonylated residue within the protein, with the fold change impact of ACSF3 KD color coded. **j.** Matrix layout for all intersections (vertical bars), sorted by size, of differentially regulated lipids of sh*Ctl* vs. sh*Acsf3* mouse liver between the lipids that loss or gain rhythmicity or with significant up-regulated (UP) or down-regulated (DOWN) mesor. Color-coded circles in the matrix indicate sets that are part of the intersection and, the top horizontal bar graph displays the metabolites number. Each category of lipids is color coded. The inset pie plot displays the proportion of lipids gaining (blue), losing (dark red) or conserving (green) rhythmicity upon ACSF3 KD or non-rhythmic lipids (white). **k.** Rhythmic lipid class and differentially regulated proteins at total and malonylated levels in triglycerides synthesis pathway (adapted from KEGG pathway database). Each dot represents a class of lipids or a lipid, with color coding representing differential levels or changes in rhythmicity. Each rectangle represents a protein and the impact of ACSF3 KD on protein fold change is color coded.

Overall, we observed that ∼60% of the metabolites were rhythmic in sh*Ctl* or sh*Acsf3* mouse liver with half remaining rhythmic upon ACSF3 KD (Fig. 6.f, Table S4). These rhythmic metabolites showed similar phase and amplitude upon ACSF3 KD (Table S4; Fig. S6a; Fig. S6b) but presented a significant global phase delay (Fig. S6c).

Among the 23% metabolites losing rhythmicity in ACSF3 KD animals, we observed a majority of fatty acids (e.g. Palmitoleic acid (C16:1), Stearic acid (C18:0), Oleic Acid (C18:1), Linolenic acid (C18:2), (Fig. 6g)) and acyl-carnitines (e.g. palmitoyl-carnitine (Fig. 6h)) included in the 23% metabolite group losing rhythmicity (Fig. 6f ; Fig. 6i Table S4). Of interest, carnitine and acetyl-carnitine (Fig. 6g) were also part of the 8% of the metabolites gaining rhythm, (Fig. 6f; Fig. 6h Table S4) while long chain acyl-carnitines such as myristoyl-carnitine (C14), and palmitoyl-carnitine (C16) were present in the group of metabolites significantly up regulated (Fig. 6h ; Fig. S6d ; Table S5).

Our proteomics data (Fig. 5 and Table S3) showed proteins implicated in lipid metabolism (ACSL1, CPT2, EHHADH, HDHA, ACAA1A and ACAT) (Fig. 6i; Fig. S6e) were significantly downregulated at ZT0 at the total and/or malonylation level upon ACSF3 KD. To get an idea about the potential impact of ACSF3 KD on fatty acid import within the mitochondria, we plotted the temporal ratio between each acyl-carnitine and its corresponding acyl-CoA and observed for four of them a significant increase upon ACSF3 KD (Fig. S6f). These observations suggest that lipid synthesis and β-oxidation diurnal regulation is controlled by ACSF3, while ACSF3 KD promotes lipogenesis.

To confirm this result, we performed lipidomic analysis on the same samples and observed that, the lipidome composition of sh*Ctl* vs sh*Acsf3* mouse liver was globally similar with a trend for more TG content in sh*Acsf3* mouse liver (Fig. S6g). Differential rhythmicity analysis using CircaCompare (see Methods; Table S5) revealed that 81% of the rhythmic lipids lost their rhythmicity upon ACSF3 KD when only 2% and 1.5% conserved or gained rhythmicity. The vast majority of these lipids were TG and diglycerides (DG) (Fig. 6j).

Our data also showed that 8% of the lipids were affected by ACSF3 KD (Fig. 6j ; Table S5). Interestingly, the majority of the upregulated lipids were TG (Fig. 6j Fig. S6h). In addition, when we looked at the level of total TG over time in sh*Ctl* vs sh*Acsf3*, we observed that the disruption of TG rhythmicity is accompanied by an increase of TG in the second part of the light period (Fig. S6i). Furthermore, proteomics data showed that PLIN3 and LDAH, two proteins implicated in liver TG storage regulation^47,48^ are significantly upregulated upon ACSF3 KD at ZT0. Overall abundance level or the malonylation level of proteins implicated in TG synthesis were not modified at ZT0. However, intermediates of TG synthesis such as DG and glycerophosphatidic acid (PA) also have their rhythmicity disturbed upon ACSF3 KD (Fig. 6k; Fig. S6i; Table S5).

Altogether these results show that ACSF3 KD disturbs diurnal lipid metabolism (Fig. S6i) and rewires hepatic metabolism eventually leading to increased lipogenesis and TG synthesis.

### Daily liver glucose and mitochondrial metabolism suggest that ACSF3 KD lead hepatic metabolism towards fuelling lipogenesis

In addition to increased DG and TG level, among the hallmarks of hepatic insulin resistance, we also observed that some ceramides level significantly increased upon ACSF3 KD (Fig. S6h). Among the 44 ceramides quantified (including ceramides, hexosylceramides and dihexosylceramides), ∼25% were rhythmic and all lost their rhythmicity (Table S5). Moreover, the global mean level of ceramides and hexosylceramides classes tended to be higher upon ACSF3 KD (Fig.7a). Targeting proteins involved in lipid and glucose metabolism, the phosphorylation of AKT is central in hepatic metabolic homeostasis. While the diurnal rhythm of AKT phosphorylation (Ser473) was preserved with a slightly delayed phase, its mean level was significantly decreased upon ACSF3 KD (Fig. 7b).

**Figure 7.**
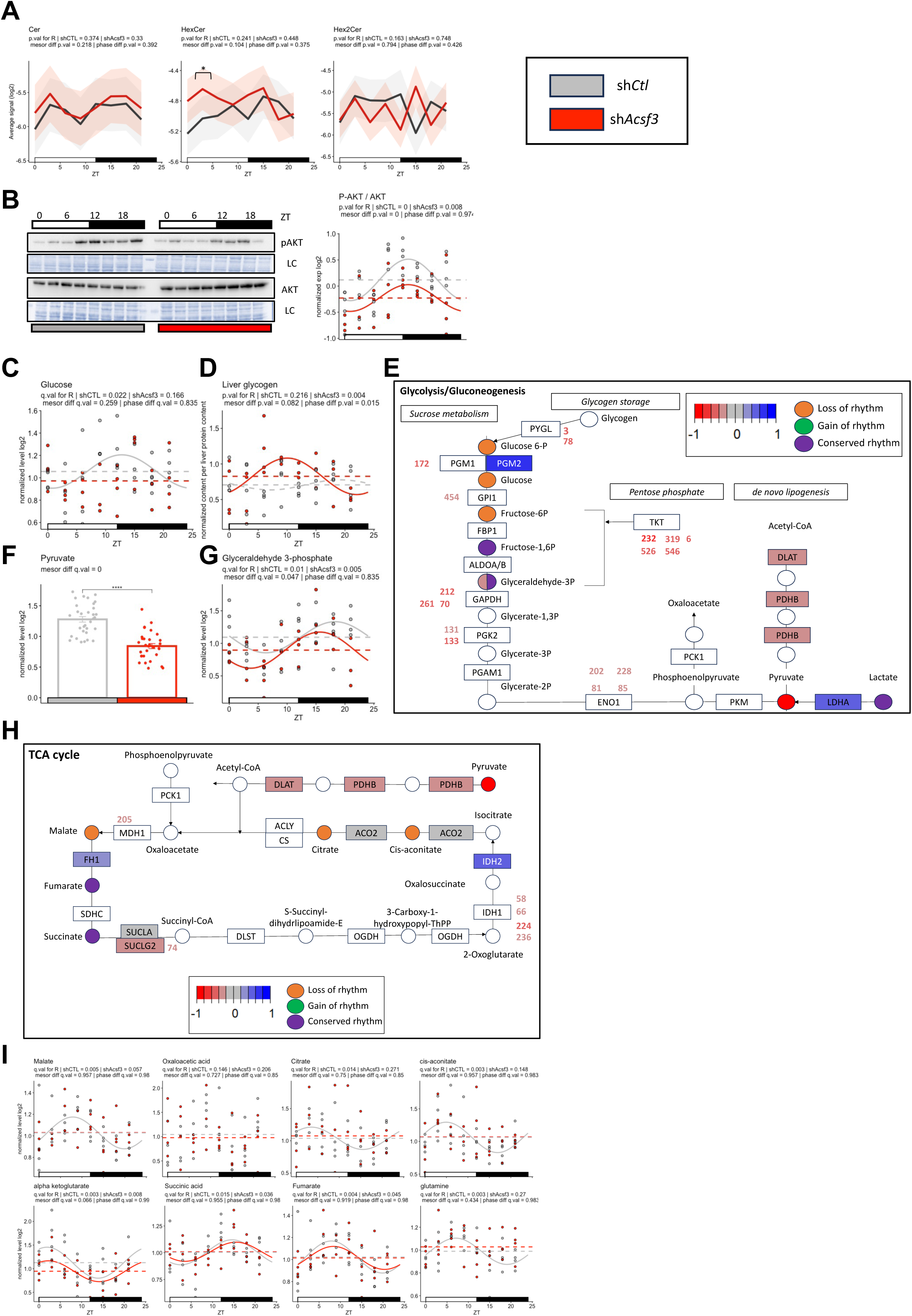
**a.** Average daily profile of ceramides (Cer), hexosylceramides (HexCer) and dihexosylceramides (Hex2Cer) in sh*Ctl* (grey) vs. sh*Acsf3* (red) mouse liver over time. Data are represented as the mean (line) ±SEM (ribbon) for each class. Rhythmicity statistics are reported. For each ZT, sh*Ctl* vs. sh*Acsf3* were compared using Tukey HSD test. **b** Western blot (WB) analysis of AKT (upper blot) and phospho-AKT (Ser 473) (lower blot) in liver TE over time in sh*Ctl* vs. sh*Acsf3* mice. The graphs display densitometry analysis of all replicates. LC: loading control. Data are presented as the ratio of P-AKT/AKT signal normalized to the global mean. Horizontal dashed lines show mesor levels for each condition. Rhythmicity statistics and fit (solid lines) are also reported. **c.** Glucose levels in TE of sh*Ctl* (grey) vs sh*Acsf3* (red) mouse liver over time. The graphs display all replicates and dashed lines correspond to mesor levels. Rhythmicity statistics and fit (solid line) are also reported. **d.** Hepatic glycogen levels in TE of sh*Ctl* (grey) vs. sh*Acsf3* (red) mouse liver over time. The graphs display all replicates and dashed horizontal lines correspond to mesor levels. Rhythmicity statistics and fit (solid lines) are also reported. **e.** Rhythmic metabolites and differentially regulated proteins at the total and malonylated levels in glycolysis/gluconeogenesis (adapted from KEGG pathway database). Each dot represents a metabolite, with color coding representing differential levels or changes in rhythmicity. Each rectangle represents a protein, and the impact of ACSF3 KD on protein fold change is color coded. The impact on malonylation levels (if present) is indicated by the presence of the position number of the malonylated residue within the protein, with the fold change impact of ACSF3 KD color coded. **f.** Pyruvate level in TE of sh*Ctl* (grey) vs. sh*Acsf3* (red) mouse liver. Mesor levels were compared with CircaCompare. **g.** Glyceraldehyde-3-phosphate level in TE of sh*Ctl* (grey) vs. sh*Acsf3* (red) mouse liver over time. The graphs display all replicates and dashed lines correspond to mesor levels. Rhythmicity statistics and fit (solid lines) are also reported. **h.** Rhythmic metabolites and differentially regulated proteins at the total and malonylated levels in the TCA cycle (adapted from KEGG pathway database). Each dot represents a metabolite with color coding representing differential levels or changes in rhythmicity. Each rectangle represents a protein and the impact of ACSF3 KD on protein fold change is color coded. The impact on malonylation levels (if present) is indicated by the presence of the position number of the malonylated residue within the protein, with the fold change impact of ACSF3 KD color coded. **i.** TCA cycle metabolite levels in TE of sh*Ctl* (grey) vs. sh*Acsf3* (red) mouse liver over time. The graphs display all replicates and dashed lines correspond to mesor levels. Rhythmicity statistics and fit (solid lines) are also reported.

Insulin resistance is associated with lower levels of mitochondrial oxidative enzymes. Hence, we looked at the global expression of protein composing mitochondrial electron transport chain complexes and observed that ACSF3 KD overall decreased Complex I, III and IV level (Fig. S7a, Table S3). The blunted rhythmicity of the ATP/ADP ratio also suggest impaired diurnal oxidative phosphorylation (Fig. S7b). To further explore this, we performed Seahorse analysis on isolated mitochondria from sh*Ctl* vs. sh*Acsf3* mouse liver. sh*Acsf3* mice exhibited lower basal respiration and oxygen consumption rates (OCR) (Fig. S7b). Although the protein level of Complex II components is not impacted, its inhibition due to the increased of malonate^49^ (Fig. 6c) could also be implicated in the observed decreased oxidative phosphorylation.

In accordance with decreased AKT phosphorylation, we observed ACSF3 KD blunted glucose diurnal rhythm (Fig.7c). Hence, we quantified glycogen content and observed that sh*Ctl* mice tended to have higher glycogen at around ZT20. Interestingly, ACSF3 KD drastically phase shifted diurnal liver hepatic glycogen peaking earlier at ZT10 (Fig. 7d). Of note, proteomics data revealed that two malonylated lysine of glycogen phosphorylase (PYGL), were downregulated (Fig. 5h; Fig. 7e; Table S3). Our results suggest that ACSF3 KD disturb the daily rhythm of glucose mobilization from liver glycogen stocks and that an alteration in PYGL malonylation is likely implicated (Fig.7e).

Although not rhythmic, the glycolysis end product pyruvate was significantly decreased upon ACSF3 KD (Fig. 7e; Fig. 7f) along with a decreased level of glyceraldehyde-3-phosphate (Fig.7e;Fig7g). Glycolysis protein levels or their malonylation levels, notably GPI1 (1 site), GAPDH (3 sites), PGK1;PGK2 (2 sites) and ENO1 (4 sites) were also decreased, so as the enzyme converting pyruvate into acetyl-CoA (DLAT) to fuel the TCA cycle (Fig. 5d; Fig. 5g; Fig. 7e; Fig. 7h). Thus, we observed the diurnal regulation of TCA cycle metabolites and observed a blunted rhythm for malate, citrate and cis-aconitate and a decreased level for alpha-ketoglutarate (Fig. 7i). No impact on diurnal regulation was observed for fumarate, succinate and oxaloacetate. Interestingly, IDH2 protein level increased and four of IDH1 malonylated lysine were downregulated suggesting an impact of ACSF3 KD on their regulation (Table S3; Fig. 5d). Of note, we also observed that glutamine, an amino acid that can mediate *de novo* lipogenesis via IDH1^50^, had its rhythmicity blunted upon ACSF3 KD (Fig. 7i), so as citrate (Fig. 7i) and this was synchronous with the observed global loss of fatty acid synthesis rhythmicity (Fig. 6).

Altogether, our observations suggest that ACSF3 KD impair diurnal liver energy metabolism. The diurnal signature of mitochondria-linked metabolites suggests a metabolic disequilibrium rewired towards increased fatty acid synthesis.

## DISCUSSION

In the present work, we show that ACSF3 diurnal expression and mitochondrial lysine-malonylation are rhythmic and synchronous, with ACSF3 peaking a bit earlier under night-restricted feeding. Previous work using mouse model of obesity showed that disturbance in hepatic mitochondrial proteins malonylation reflect liver ACSF3 protein abundance^29^. Although ACSF3 and lysine-malonylation rhythmicity is disturbed in DIO, they maintain synchrony in all conditions studied confirming the link between ACSF3 expression and the regulation of mitochondrial protein malonylation. Taking advantage of previously published proteomics^5,12^ and transcriptomics data^37–39,43^, our analysis show that the molecular circadian clock does not appears to be involved in ACSF3 diurnal expression rhythmicity. In fact, our study shows that feeding schedule impact ACSF3 expression as well as mitochondrial malonylation and is rather the main driver of their diurnal rhythmicity.

Upon mistimed feeding or DIO, the misalignment between timing cues such as glucocorticoids and insulin signalling affects the robustness and disturbs circadian rhythms in peripheral organs such as the liver^51,52^. A transcriptomic study from Fougeray *et al.* indicate that the impact of dRF on hepatic *Acsf3* expression relies on a functional insulin signalling^53^. Quagliarini *et al.* showed that DIO expands the Glucocorticoid Receptor cistrome. In particular, their data reveal that a region near the transcription start site (TSS) of the *Acsf3* promoter is included in the Glucocorticoid Receptor cistrome upon HFD at ZT12^43^. The same observations were made for *Sirt5* (intergenic) and *Mlycd* (near TSS) genes, suggesting glucocorticoid signalling also regulates diurnal mitochondrial malonylation regulators upon DIO^43^. Thus, glucocorticoid signalling might play a role in the observed phase shift of *Acsf3* expression. Altogether, these observations suggest that combined disturbance within insulin and glucocorticoid signalling pathways explain the differential regulation of *Acsf3* expression observed upon dRF and HFD. Overall, these signalling pathways would be implicated in driving rhythmic ACSF3 expression and mitochondrial lysine-malonylation.

In a genetic mouse model of obesity, ACSF3 expression as well as mitochondrial malonylation are differentially deregulated in the liver (upregulated) vs. other peripheral organs such as heart, brown adipose tissue (down-regulated) or kidney (not affected)^29,54^. Hence, we decided to specifically study the impact of hepatic ACSF3 KD that did not protect nor sensitize mice to metabolic abnormalities upon DIO which is reminiscent of *Sirt5* KO mice phenotype^45^. However, while *Sirt5* KO mice are slightly more insulin sensitive and present increased hepatic lysine-malonylation level^24,45^, the opposite is observed upon ACSF3 KD (*i.e.* impaired insulin sensitivity coupled with decreased hepatic lysine-malonylation) suggesting lysine-malonylation events could explain the afore described opposite phenotypes. Although the metabolic pathways enriched in SIRT5-regulated vs. ACSF3 impacted malonylome are common, our data suggest poor functional overlap.

In fact, we showed that ACSF3 KD impairs the diurnal regulation of hepatic glycogen stocks and rewires hepatic metabolism to TG synthesis. The diminution of insulin sensitivity, coupled with the increased glucose tolerance, suggests a selective insulin resistant state where the liver is unable to regulate properly diurnal glucose production and promotes *de facto de novo* lipogenesis and TG synthesis. We have not specifically directly tested the functionality of the quantified malonylated lysine (365 in the current study). However, the fact that within the deregulated pathways we observed key regulatory proteins with significant differentially regulated malonylated lysines (e.g. 2 sites within PYGL for glycogen content regulation, or 4 in IDH1) suggests they could be functionally directly implicated in the observed metabolic phenotype like already shown for several examples of malonylated lysines^55–57^. On the other hand, among the differentially impacted malonylated lysine by ACSF3 KD, most if not all can also be modified by multiple other type of PTMs such as ubiquitination or other acylations like acetylation, glutarylation or succinylation with potential consequences on their stability or functionality^24,26,58,59^. IDH1 is of interest since three of its malonylated-lysine that are down-regulated can also be ubiquitinated. Thus, ACSF3 KD driven malonylation decreased could favour IDH1 lysine-ubiquitination favoring its unstability. Besides lysine-ubiquitination, the example of lysine-succinylation is also of interest. Indeed, increased lysine-succinylation is also associated with increase acyl-carnitines level and modified β-oxidation^58^. Thus, malonylation decreased upon ACSF3 KD could also favor succinylation occurrence, leading to part of the functional metabolic consequences observed.

Results from our mice work are also aligned with previous work studying metabolism of fibroblasts from patients with homozygous mutations of ACSF3 genes resulting in a loss of ACSF3 function. Whebe *et al.* observed increased level of taurine, decreased level of glutamine, threonine, ceramides and increased level of TG, while we show an impact on their diurnal rhythmicity and/or mean level upon ACSF3 KD^28^. The latter work as the work of Bowman *et al.* also observed a significant decrease in oxidative phosphorylation upon ACSF3 loss of function^28^ or ACSF3 KO^29^. We show a decreased of oxidative phosphorylation as one of the functional consequences of ACSF3 KD. This could be linked to global impact on the expression of protein composing mitochondrial electron transport chain complexes I, III or IV. ACSF3 potential role in finely tuning the level of important lipids required for proper electron transport chain assembly such as cardiolipins is also not to be excluded^60,61^. Although the protein level of its components is not impacted, Complex II can also be inhibited by the global increase of malonate level upon ACSF3 KD^49^. Our study suggests the ACSF3-controlled rhythmicity of malonate could be an additional mechanism in driving the diurnal rhythmicity of oxidative phosphorylation.

Our proteomic study shows that ACSF3 KD differentially regulates the abundances of proteins at the total level without impacting their malonylation state. This suggests an effect via other PTMs but also via a global impact of ACSF3 KD on gene expression. In fact, the human proteome atlas database indicates ACSF3 also localizes in the nucleus^62^. Via producing malonyl-CoA, ACSF3 could locally influence the malonylation of histones with functional consequences on chromatin function and gene expression^26,63–65^.

In conclusion, we showed ACSF3 and its mediated mitochondrial malonylation are rhythmic and controlled by feeding schedule. While hepatic ACSF3 does not modify DIO metabolic abnormalities, our multi-omics approach established a role for ACSF3 in diurnal regulation of liver glycogen storage, *de novo* lipogenesis and TG synthesis.

## AUTHOR CONTRIBUTIONS

E.L. & D.M. conceptualized the study.

E.L., C.B., F.A., Y.F. & D.M. carried out the investigation.

A.T., V.P., A.R.O., M.C. & Y.C. performed the proteomics experiments.

M.G., M.C. & I.N. performed the metabolomics experiments.

C.C. & M.C. performed the lipidomics experiments.

E.L & D.M. curated the data.

E.L & D.M. carried out the data analysis.

E.L & D.M. carried out the data visualization.

D.J., B.C. & D.M. obtained the funding.

E.L. & D.M. wrote the original draft of the manuscript.

E.L., F.A., I.N., M.C. Y.C., D.J., B.C. & D.M. edited and reviewed the manuscript.

## ACKNOWLEDGMENTS

This research was supported by the French National Research Agency under the “Programme Investissement d’Avenir” (NExT (I-SITE); ANR-16-IDEX-0007) and ANR R20109NN. Additional funding was provided by the French Regional Council of Pays de la Loire through the PULSAR “Académie des Jeunes Chercheurs en Pays de la Loire,” the “NSFA Nouvelle Société Francophone d’Athérosclérose,” and the “Fondation Genavie.” The proteomic experiments were partially supported by the Agence Nationale de la Recherche under the projects ProFI (Proteomics French Infrastructure, ANR-10-INBS-08) and GRAL, a program from the Chemistry Biology Health (CBH) Graduate School of the University Grenoble Alpes (ANR-17-EURE-0003).

The authors thank Frédéric Gachon and Benjamin Weger for critical reading of the manuscript.

## METHODS

### Animal experiments

Animal studies were conducted in accordance with the regulations of the veterinary office of the region Pays de la Loire (Apafis #26868 and #37476 in compliance with directive 2010/63/EU). SILAC mice generated according to standard procedure^66^ has been previously described^5^.

Food access was controlled via programmable automatic feeders (SmartWaiter, Cibertec, Madrid) and mouse were single caged upon food intake monitoring. For all experiments, if not differently indicated, mice had free access to food and water in 12 hours light/12 hours dark cycles under standard animal housing conditions.

#### High fat diet experiment

At 12 weeks mice were fed with hight fat diet (60% fat, D12492, Research Diet) or standard chow diet (SAFE A04 Rosenberg, Germany) *ad libitum* during 12 weeks.

#### Acsf3 hepatic knock down

experiments were performed on mice fed a standard or HFD diet. Mice were injected retro-orbitally with AAV8-sh *Acsf3* and control vectors (Vector Builder).

#### Metabolic phenotyping

at the indicated weeks during experiments, a glycemia follow up was performed at ZT0. Mice were fasted during 6 hours (from ZT0 to ZT6) before intraperitoneal insulin (ITT) and glucose (GTT) tolerance tests were performed. For the GTT, 1.5 g/kg of glucose was intraperitoneally injected and glycemia measure were performed at 20, 40, 60, 90 and 120-min post-injection. For the ITT 0.75 U/Kg of insulin was intraperitoneally injected followed by glycemia measured at 15,30,60,90 and 120 min post -injection. Data were analysed according to the following guidelines^67,68^.

### Isolation of liver mitochondria

Liver mitochondria were isolated by differential centrifugation as previously described^12^, with minor modifications. Detailed experimental procedure is described in Supplemental Information.

### Bioenergetics analyses

The oxygen consumption rate was determined on a Seahorse XF96 HS Mini Analyzer. For mitochondrial coupling assays, isolated mitochondria were incubated in a mitochondrial assay buffer (70 mM sucrose, 220 mM mannitol, 10 mM KH2PO4, 5 mM MgCl2, 2 mM HEPES, 1 mM EGTA, and 0.2% fatty acid free BSA, pH 7.2) with 10 mM succinate. Sequential addition of 4 mM ADP, 2 µg/mL oligomycin, 4 µM FCCP, and 4 µM antimycin A were used to determine state 3 and state 4 respiration.

### Total and mitochondrial protein extraction

Livers protein total extracts (TE) or mitochondria protein extracts (ME) from liver mitochondria were prepared as previously described^32^. Detailed experimental procedure is described in Supplemental Information. **Western Blotting**

Western blotting was performed according to standard procedures. Densitometry analyses of the blots were performed using the ImageJ software. Naphtol blue black staining of the membranes was used as a loading control and served as references for normalisation of the quantified values. A list of antibiodies used in this study is available in Supplemental Information.

### Total Liver vs. mitochondrial liver proteomic data analysis

Proteomics data of liver total extract from WT, *Cry1/2* KO, *Bmal1* WT and *Bmal1* KO (TE) were from (PXD001211)^32^. Proteomics data from mitochondria extract (ME) were from (PXD001732)^12^. Rhythmicity analysis was assessed using CircaCompare^33^. Mean level difference of proteins from TE in *Bmal1* KO, *Cry1/Cry2* KO vs. their corresponding littermates^5^ was assessed using ANOVA (FDR <0.05).The detailed procedure can be found in Supplemental Information.

### Transcriptomics data analysis

Rhythmicity of RNAseq data from *Bmal1* KO or *Cry1/2* KO or their corresponding littermates fed chow diet under night restricted feeding regimen (GSE135898)^37^, of mice fed chow diet *Ad libitum* or night restricted feeding regimen (GSE118967)^38^, of mice fed chow diet *Ad libitum* or day restricted feeding regimen (GSE159135) ^39^ or of mice on chow diet versus HFD (GSE108688)^43^ was assessed using CircaCompare^33^. The resulting *p-values* were used to estimate FDR by the Benjamini-Hochberg method^69^. Statistical significance was set to FDR = 0.05. (Table S2).

### Functional annotation (GO TERM), KEGG pathway and enrichment analysis

Enrichment for Mitocarta 3.0 pathway^34^ were calculated using a Fisher’s exact test. Gene ontology (GO)^70^ analysis was performed with Pantherdb^71^. Enrichment test was performed for Cellular component (CC_all). The KEGG pathway^72^ enrichment for quantified proteins was performed using cluster profiler^73^ package. Top 10 enriched categories with adjusted p-values (Benjamini–Hochberg) smaller than 0.05 were reported. The gene universe was defined as all quantified proteins.

### Proteomic analysis

#### Sample preparation for proteomic analysis

Proteins from mouse hepatic TE from non-SILAC mice (*i.e.* 5 shCTL mice and 5 sh*Acsf3* mice) were collected at ZT0 and prepared as in^74^. The detailed procedure can be found in Supplemental Information. (Table S3).

#### MS-based proteomic analysis of proteome and Malonyl-proteome

MS/MS based SILAC proteomic analysis is mostly done as previously described^74^. Detailed experimental procedure is described in Supplemental Information.

#### Untargeted Metabolomic analyses

Metabolomics analyses were performed on liver and isolated hepatic mitochondria and done as previously described^75^. Rhythmicity analysis was assessed using CircaCompare^33^.The resulting *p-values* of all metabolites were used to estimate FDR by the Benjamini-Hochberg method^69^. Detailed experimental procedure is described in Supplemental Information. (Table S4).

#### Quantification of malonic acid and methyl-malonic acid

Detailed experimental procedure is described in Supplemental Information.

#### Mass spectrometry-based lipidomics

Non targeted lipidomic analysis is mostly done as previously described^76^ and the identification as previously described^77,78^. Rhythmicity analysis was assessed using CircaCompare^33^.The resulting *p-values* of all lipids were used to estimate FDR by the Benjamini-Hochberg method^69^. The threshold for significant rhythmicity was set to p.value <0.05 which correspond to a FDR = 0.17 according to previous work from Aviram *et al.* 2016^79^. Detailed experimental procedure is described in Supplemental Information.

#### Liver glycogen content measurements

Liver glycogen content was determined according to the manufacturer’s protocol (MAK016 Glycogen assay kit, Sigma-Aldrich).

## LIST OF TABLES

**TableS1: Proteomics Data Analysis**

**Datasheet1:** CircaCompare analysis of the hepatic proteome and the mitochondrial proteome. (PXD001211and PXD001732).

**TableS2: Transcriptomics Data Analysis**

**Datasheet1:** CircaCompare analysis of RNA-Seq results from animal models with a genetically disrupted circadian clock under a *night-restricted* feeding regimen (*Bmal1* WT vs *Bmal1* KO) (GSE135898).

**Datasheet2:** CircaCompare analysis of RNA-Seq results from animal models with a genetically disrupted circadian clock under a *night-restricted* feeding regimen (*Cry1/Cry2* WT vs *Cry1/Cry2* KO) (GSE135898).

**Datasheet3:** CircaCompare analysis of RNA-Seq results from animals fed *ad libitum* or under a *night-restricted* feeding regimen (GSE118967).

**Datasheet4:** CircaCompare analysis of RNA-Seq results from animals fed *ad libitum* or under a *day-restricted* feeding regimen (GSE159135).

**Datasheet4:** CircaCompare analysis of RNA-Seq results from animals fed *ad libitum* on a CD or HFD regimen (GSE108688).

**TableS3: Proteomics Data Analysis**

**Datasheet1:** Quantification of the hepatic malonyl-proteome in animal models with a hepatic knock down of *Acsf3* under an *ad libitum* feeding regimen (sh*Ctl* vs sh*Acsf3*).

**Datasheet2:** Quantification of the hepatic proteome in animal models with a hepatic knock down of *Acsf3* under *an ad libitum* feeding regimen (sh*Ctl* vs sh*Acsf3*).

**TableS4: Metabolomics Data Analysis**

**Datasheet1** CircaCompare analysis of hepatic metabolomics results from animal models with a hepatic knock down of *Acsf3* under an *ad libitum* feeding regimen (sh*Ctl* vs sh*Acsf3*).

**TableS5: Lipidomics Data Analysis**

**Datasheet1** CircaCompare analysis of hepatic lipidomics results from animal models with hepatic knockdown of *Acsf3* under an *ad libitum* feeding regimen (sh*Ctl* vs sh*Acsf3*).

## SUPPLEMENTAL FIGURE LEGEND

**Figure S1.**
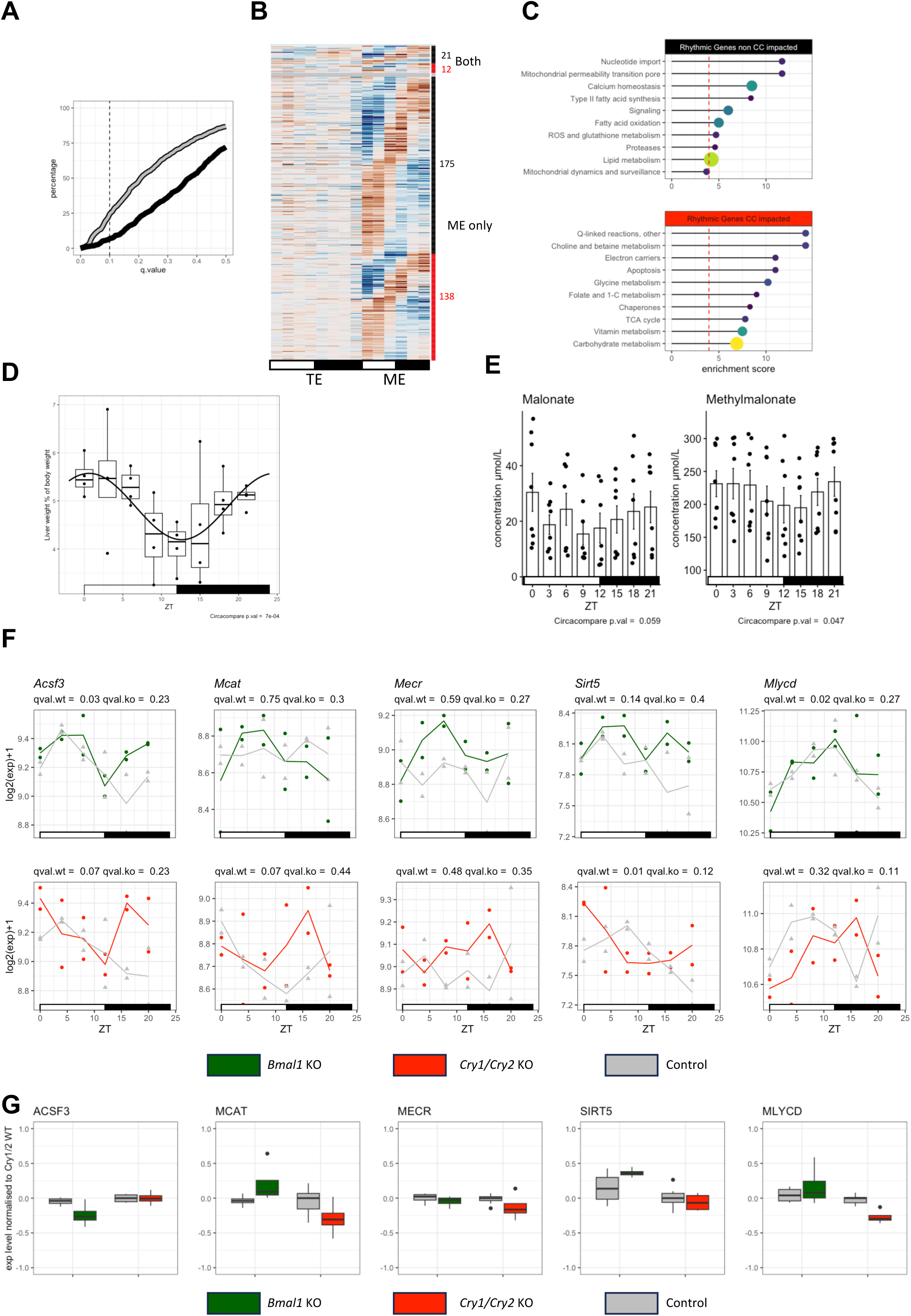
**a.** Percentage of rhythmic proteins in liver TE (black) or ME (grey) over a range of FDR up to 0.5. Vertical dashed line shows FDR=0.1. **b.** Heatmaps of protein expression in mouse liver TE (left) and ME (right) of all rhythmic proteins in ME detected with CircaCompare (FDR < 0.1). Data are clustered by rhythmicity and by circadian clock deficiency impact on mean expression (see Methods) in TE (right bar). **c.** Top 10 enriched Mitocarta 3.0 pathways in proteins with rhythmic expression only in ME with mean expression levels perturbed (lower panel) or not perturbed (upper panel) in circadian clock deficient animals. Vertical dashed line shows global median enrichment score. **d.** Liver weight over time. Black line corresponds to the rhythmicity fit. Caption indicates p.value for rhythmicity. **e.** Liver malonate (left panel) and methylmalonate (right panel) levels over time (Data presented as mean ± SEM). Caption indicates p.value for rhythmicity. **f.** Gene expression of *Acsf3*, *Mcat*, *Mecr*, *Sirt5* and *Mlycd* over time in WT or circadian clock deficient mouse liver. Upper and lower panels are respectively *Bmal1* KO (green) and *Cry1/2 KO* (red) and their corresponding controls (grey). The graphs display all replicates and lines correspond to mean level over time. Statistics for rhythmicity are also reported. **g.** Protein expression of ACSF3, MCAT, MECR, SIRT5 and MLYCD in WT or circadian clock deficient mouse liver. In each panel, the left pair and right pair of boxplots are respectively *Bmal1* KO (green) and *Cry1/2 KO* (red) and their corresponding controls (grey).

**Figure S2.**
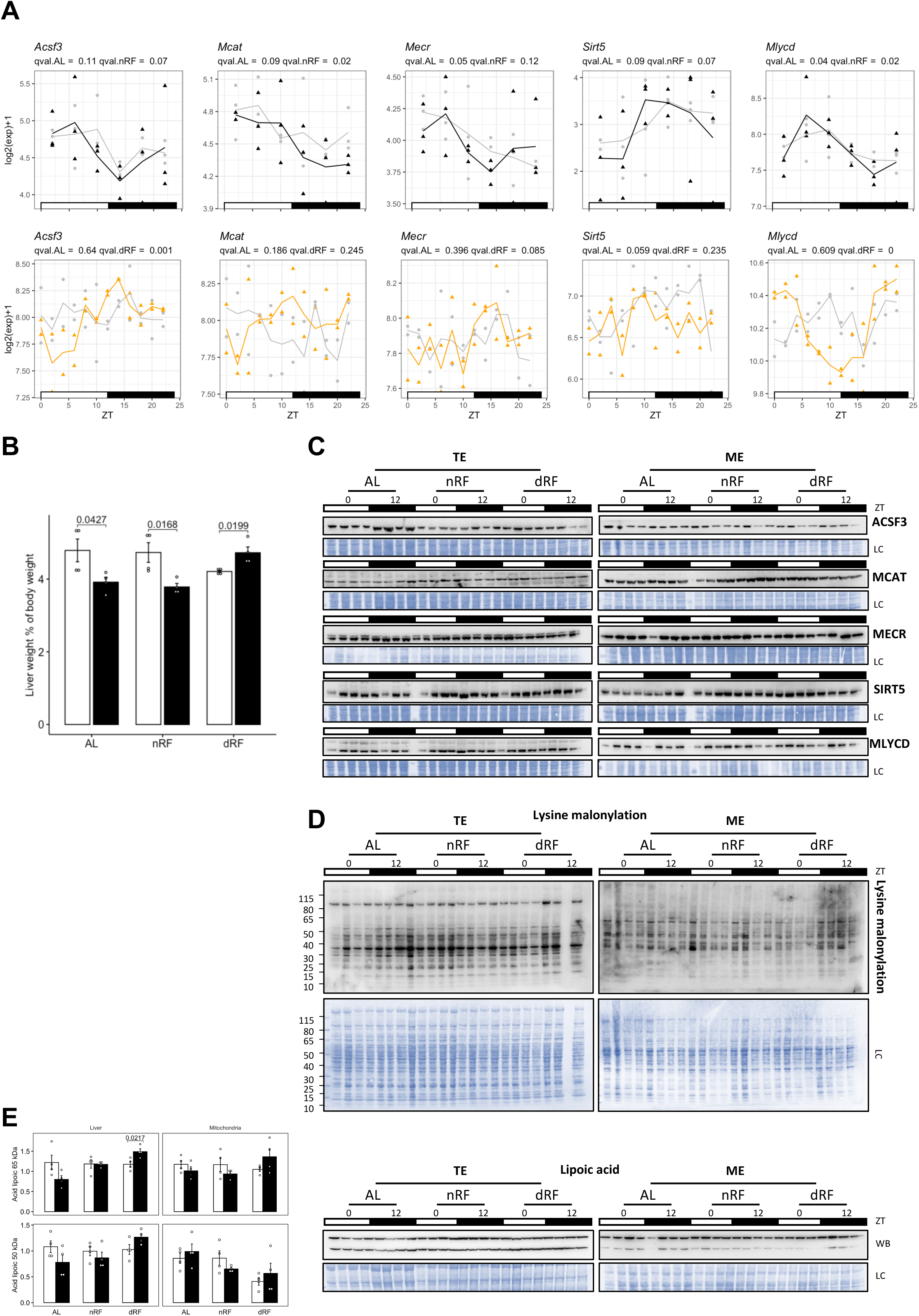
**a.** Gene expression of *Acsf3*, *Mcat*, *Mecr*, *Sirt5* and *Mlycd* over time in mice fed AL (grey upper graphs) vs. nRF (black upper graphs) or AL (grey lower graphs) vs. dRF (orange lower graphs). The graphs display all replicates and lines correspond to mean level over time. Rhythmicity statistics are also reported. **b.** Liver weight over time (n=4 replicates). ZT0 vs. ZT12 were compared using Tukey HSD test. **c-d.** Western blot (WB) analysis of ACSF3, MCAT, MECR, SIRT5, MLYCD **(c)**, and lysine malonylation **(d)** in total extracts (TE) and mitochondrial extracts (ME) (n=4 replicates) over time. LC: loading control. **e.** Western blot (WB) analysis of lipoic acid in total extracts (TE) and mitochondrial extracts (ME) (n=4 replicates) over time. The graphs display densitometry analysis of all replicates. The upper and lower graphs represent respectively the quantification of the 65 and 50 kDa bands. LC: loading control. For each extract, ZT0 vs. ZT12 were compared using Tukey HSD test.

**Figure S3.**
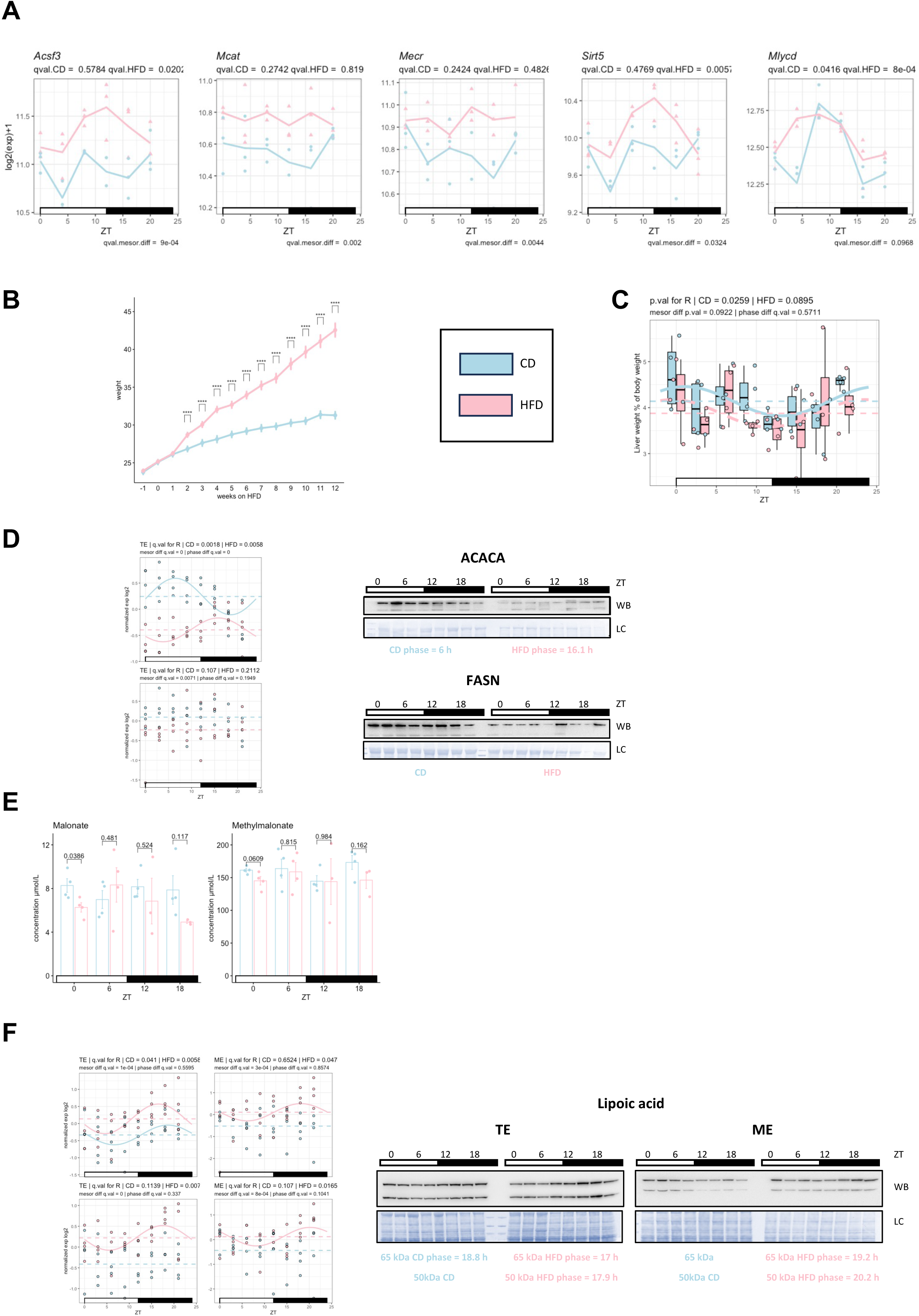
**a.** Gene expression of *Acsf3*, *Mcat*, *Mecr*, *Sirt5* and *Mlycd* over time in mice fed CD (light blue) vs. HFD (pink) over time. The graphs display all replicates and lines correspond to mean level over time. Statistics for rhythmicity are also reported. **b.** Mouse body weight in g (n=32/condition). **c.** Liver weight over time (n=4 replicates). Horizontal dashed lines show mesor level for each diet. Rhythmicity statistics and fit (lines) are also reported. **d.** Western blot (WB) analysis of ACACA and FASN in total extracts (TE) over time upon CD (light blue) or HFD (pink). The graphs display densitometry analysis of all replicates. LC: loading control. Data are normalized to the global mean in each extract. Horizontal dashed lines show mesor level for each diet. Rhythmicity statistics and fit (lines displayed if q.val < 0.05) are also reported. **e.** Liver malonate (left panel) and methylmalonate (right panel) levels over time (Data presented as mean ± SEM). For each time point, CD vs. HFD were compared using Tukey HSD test. **f.** Western blot (WB) analysis of lipoic acid in total extracts (TE) and mitochondrial extracts (ME) (n=4 replicates) over time upon CD (light blue) or HFD (pink). The graphs display densitometry analysis of all replicates. LC: loading control. Data are normalized to the global mean in each extract. Horizontal dashed lines show mesor level for each diet. Rhythmicity statistics and fit (lines displayed if q.val < 0.05) are also reported.

**Figure S4.**
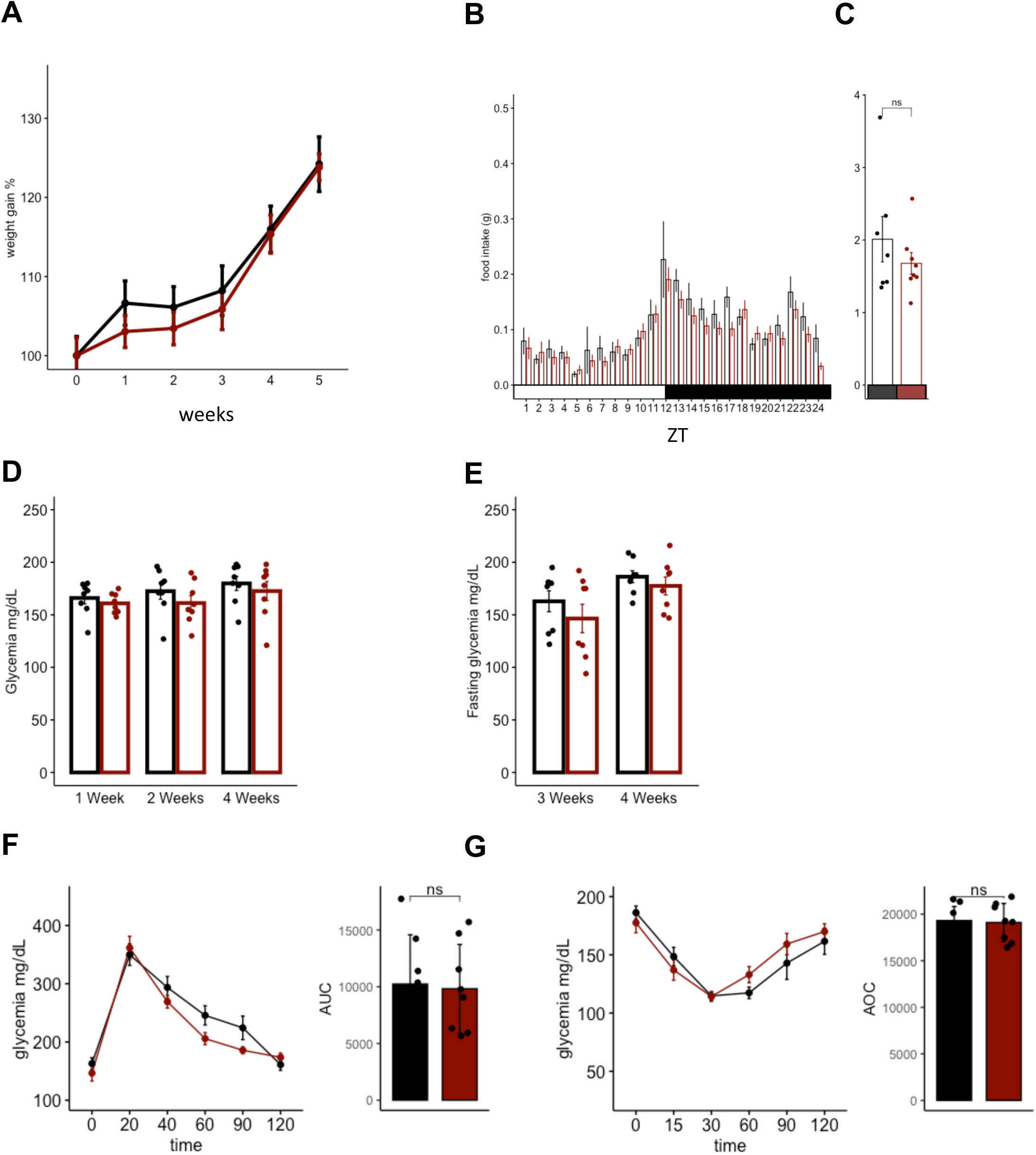
**a-g.** Mouse body weight gain (n=16/condition) **(a);** diurnal food intake **(b);** and total daily food intake **(c);** glycemia at ZT0 at 1, 2 and 4 weeks **(d);** fasting glycemia after 6 hours of fasting starting at ZT0 at 3 and 4 weeks **(e);** GTT (left) and the corresponding AUC above baseline (right) **(f);** ITT (left) and the corresponding AOC under baseline (right) **(g)** in sh*Ctl* (black) vs. sh*Acsf3* (dark red) mice upon HFD.

**Figure S5.**
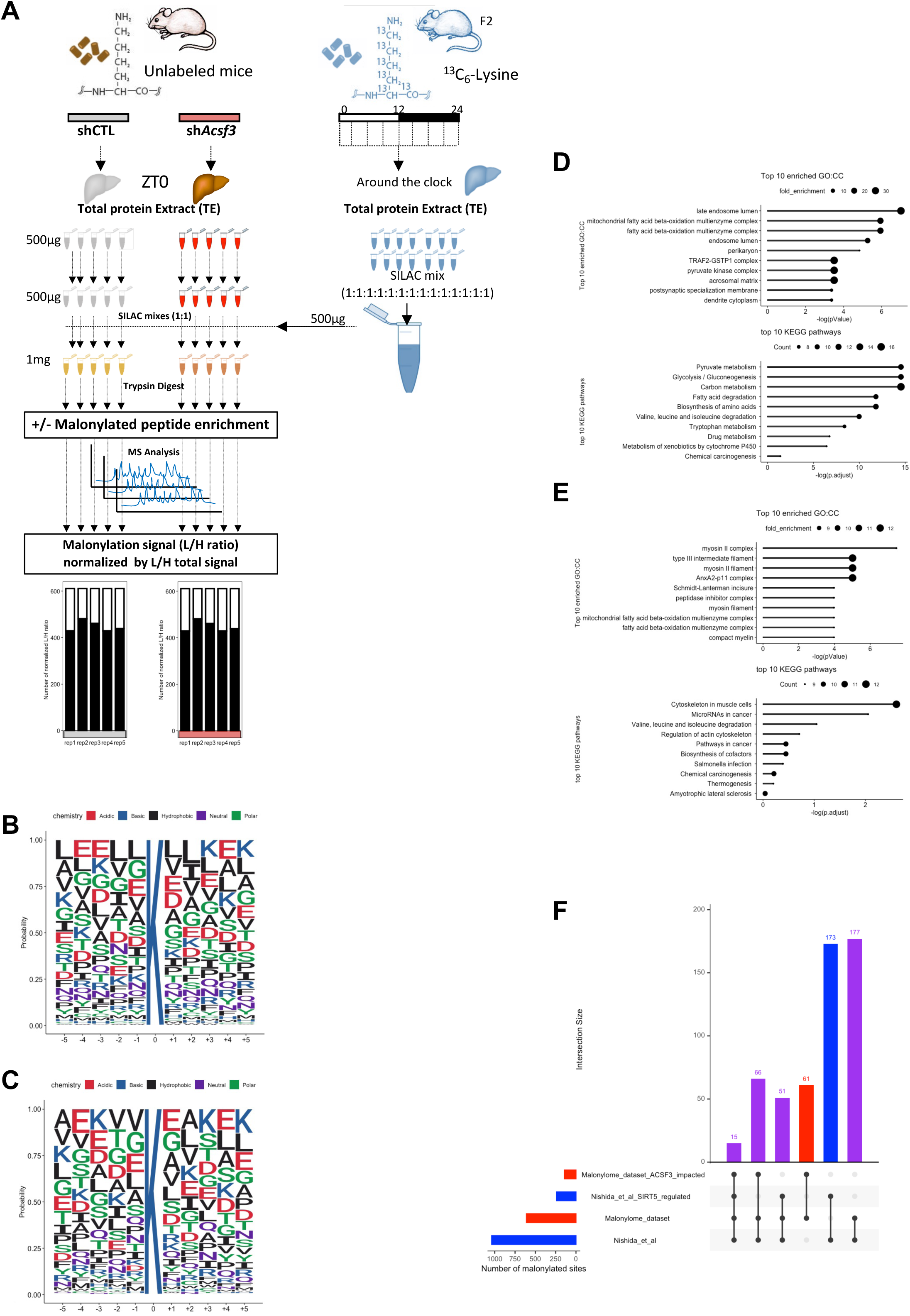
**a.** Workflow of the SILAC-based MS analysis of malonylated proteins from total mouse liver extracts (TE). The lower graph represents the number of malonylation sites for each time point. **b-c.** Consensus sequence logo plots centered on the lysine of all identified malonylation sites **(b)** or all identified SIRT5-modulated malonylation sites (±5 amino acids) **(c).** Data are from Nishida *et al.*^24^. **d-e.** Top 10 enriched GO:CC (upper graphs) and top 10 KEGG pathways (lower graphs) in all protein identified with ACSF3 KD-modulated malonylation sites **(d)** or in all protein with expression modified by ACSF3 KD **(e)**. **f-** Number of the shared differentially regulated malonylated sites between our dataset and Nishida *et al.*^24^(intersection size in purple) between all the data and the differentially regulated/impacted malonylated sites in SIRT5 KO and upon ACSF3 KD.

**Figure S6:**
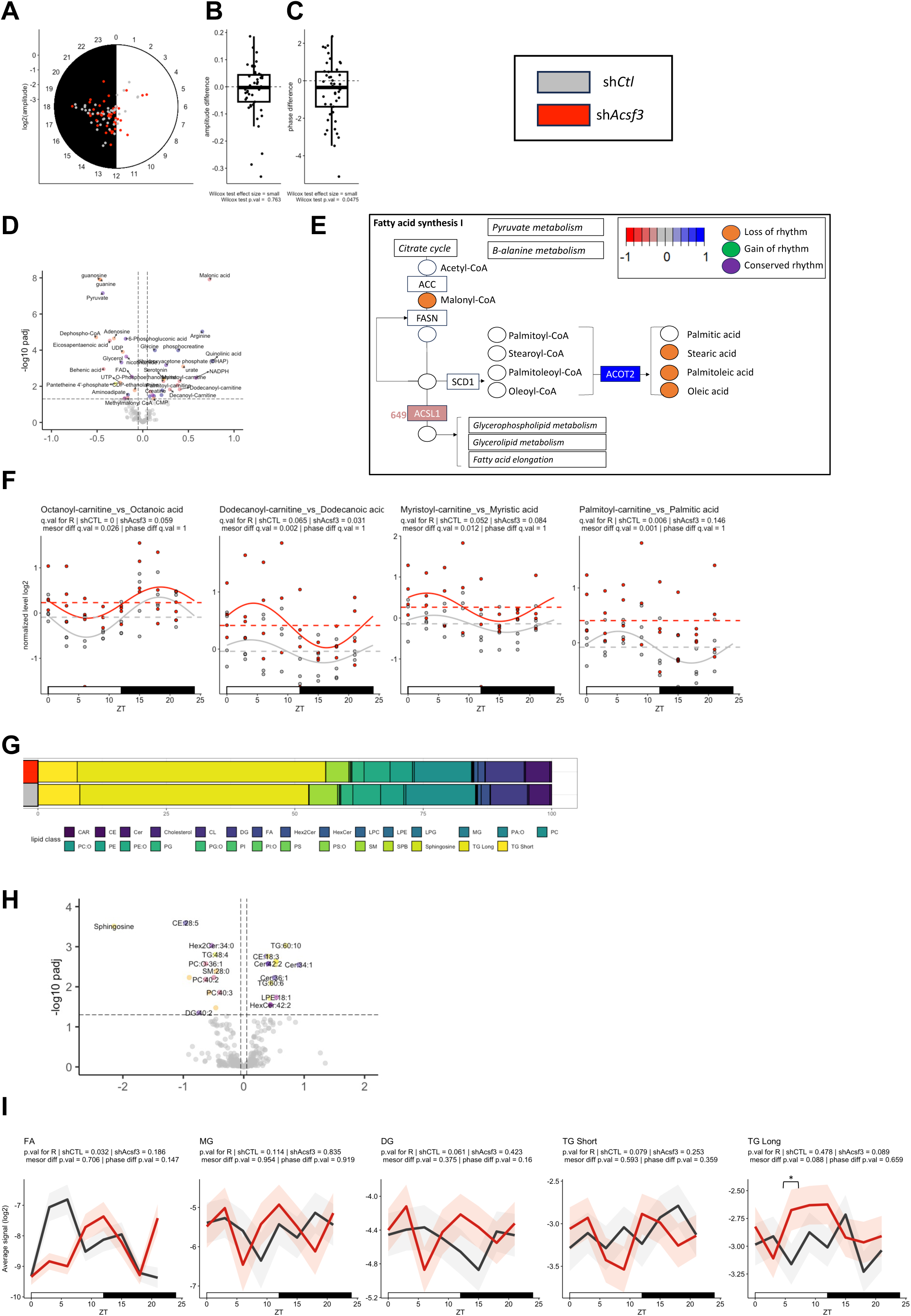
**a-c.** Peak phase (position on the circle) and amplitude (distance on the circle from the center) **(a)** amplitude **(b)** and phase difference **(c)** of metabolites conserving rhythmicity upon ACSF3 KD. **d.** Fold change imprinted by ACSF3 KD on the liver metabolome. The mesor difference is displayed. **e.** Rhythmic metabolites and differentially regulated proteins at total and malonylated levels in fatty acid synthesis (adapted from KEGG pathway database). Each dot represents a class of lipids, with color coding representing differential levels or changes in rhythmicity. Each rectangle represents a protein and the impact of ACSF3 KD on protein fold change is color coded. The impact on malonylation levels (if present) is indicated by the presence of the position number of the malonylated residue within the protein, with the fold change impact of ACSF3 KD color coded. **f.** Ratio between fatty acids vs. their corresponding acyl-carnitine levels in TE of sh*Ctl* (grey) vs sh*Acsf3* (red) mouse liver over time. The graphs display all replicates and dashed lines correspond to mesor level. Rhythmicity statistics and fit (solid lines displayed if q.val < 0.05) are also reported. **g.** Lipidome class composition of sh*Ctl* (grey) vs. sh*Acsf3* (red) mouse liver. **h.** Fold change imprinted by ACSF3 KD on the liver lipidome. The mesor difference is displayed. **i.** Average daily profile of fatty acyls (FA), monoacylglycerols (MG), diacylglycerols (DG), triacylglycerols short (TG short) and long (TG short) in sh*Ctl* (grey) vs. sh*Acsf3* (red) mouse liver over time. Data are represented as mean (line) ±SEM (ribbon) of each class. Rhythmicity statistics are also reported. For each ZT, sh*Ctl* vs. sh*Acsf3* were compared using Tukey HSD test.

**Figure S7.**
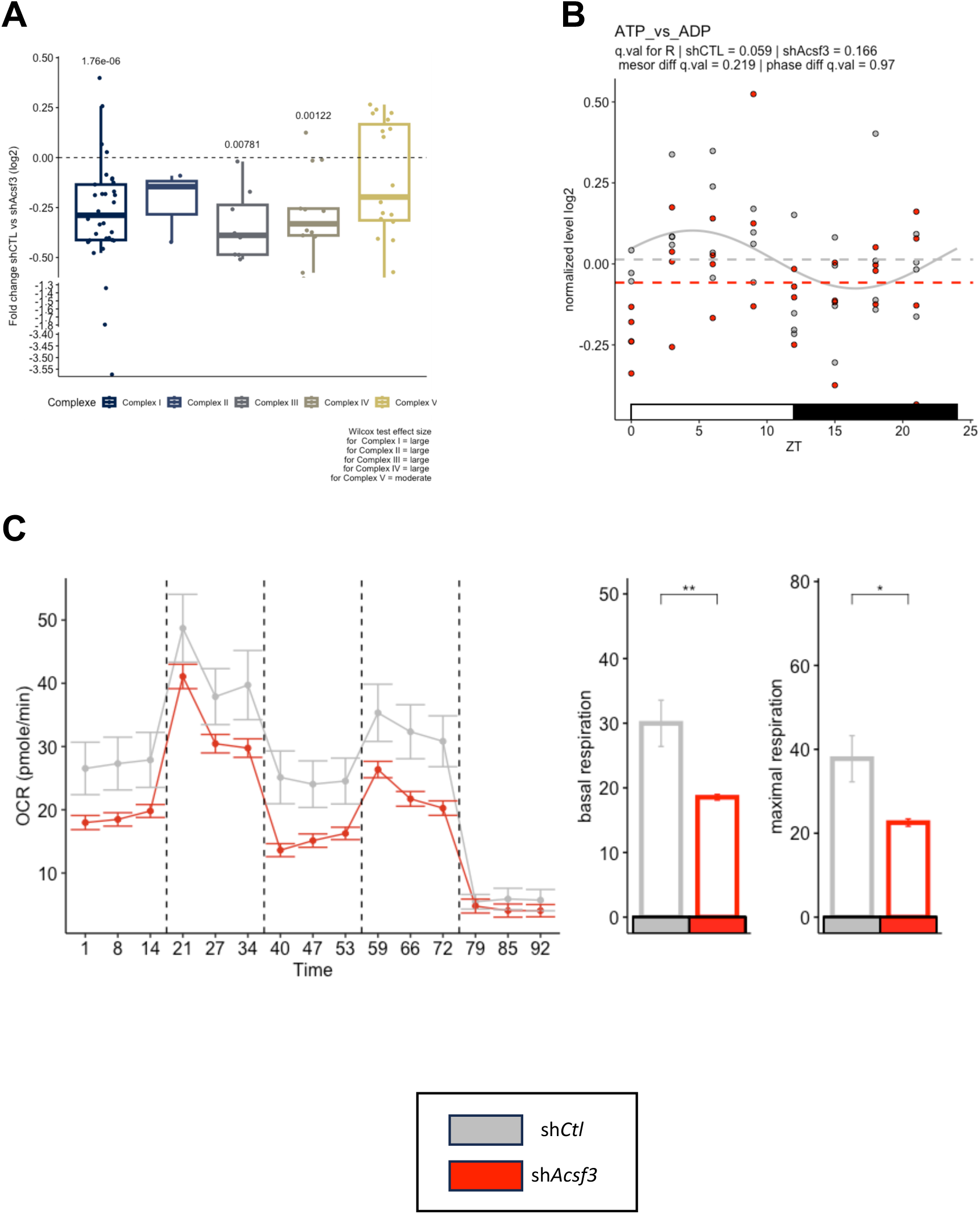
**a.** Differential impact of ACSF3 KD on expression levels of oxidative phosphorylation protein complexes components. Wilcox-test statistics and Wilcox-test effect size are reported. **b.** ATP vs. ADP levels in TE of sh*Ctl* (grey) vs. sh*Acsf3* (red) mouse liver over time. The graphs display all replicates and dashed lines correspond to mesor level. Rhythmicity statistics and fit (solid line) are also reported. **c.** Seahorse analysis on isolated mitochondria at ZT0 (n>5/condition) from sh*Ctl* (grey) vs. sh*Acsf3* (red) mouse liver. Each vertical dashed line refers to the time course of adding ADP, oligomycin, FCCP, and antimycin A respectively.

